# The apical annuli of *Toxoplasma gondii* are composed of coiled-coil and signaling proteins embedded in the IMC sutures

**DOI:** 10.1101/711432

**Authors:** Klemens Engelberg, Chun-Ti Chen, Tyler Bechtel, Victoria Sánchez Guzmán, Allison A. Drozda, Suyog Chavan, Eranthie Weerapana, Marc-Jan Gubbels

**Affiliations:** Department of Biology, Boston College, Chestnut Hill, Massachusetts, USA; Department of Chemistry, Boston College, Chestnut Hill, Massachusetts, USA; University of Puerto Rico, San Juan, Puerto Rico, USA

## Abstract

The apical annuli are among the most intriguing and understudied structures in the cytoskeleton of the apicomplexan parasite *Toxoplasma gondii*. We mapped the proteome of the annuli in *Toxoplasma* by reciprocal proximity biotinylation (BioID), and validated five apical annuli proteins (AAP1-5), Centrin2 and a methyltransferase (AAMT). Moreover, Inner Membrane Complex (IMC) suture proteins connecting the alveolar vesicles were also detected and support annuli residence within the sutures. Super-resolution microscopy (SR-SIM) identified a concentric organization comprising four rings with diameters ranging from 200-400 nm. The high prevalence of domain signatures shared with centrosomal proteins in the AAPs together with Centrin2 suggest that the annuli are related and/or derived from the centrosomes. Phylogenetic analysis revealed the AAPs are conserved narrowly in Coccidian, apicomplexan parasites that multiply by an internal budding mechanism. This suggests a role in replication, for example, to provide pores in the mother IMC permitting exchange of building blocks and waste products. However, presence of multiple signaling domains and proteins are suggestive of additional functions. Knockout of AAP4, the most conserved compound forming the largest ring-like structure, modestly decreased parasite fitness *in vitro* but had no significant impact on acute virulence *in vivo*. In conclusion, the apical annuli are composed of coiled-coil and signaling proteins assembled in a pore-like structure crossing the IMC barrier maintained during internal budding.

## Introduction

Apicomplexan parasites are obligate intracellular parasites posing significant burdens on human health and livestock. These parasites have an elaborate cytoskeleton that plays key roles in their pathogenesis by facilitating invasion of their host cells as well as driving cell division. *Toxoplasma gondii,* which causes opportunistic and congenital diseases in humans, serves as an apicomplexan model for these aspects of pathogenesis (Gubbels & Morrissette, 2013, Anderson-White *et al.*, 2012, Chen *et al.*, 2015, Chen *et al.*, 2016). *Toxoplasma* develops by an internal budding process producing either two daughters in the intermediate host (endodyogeny in tachyzoites and bradyzoites), or up to eight daughters in the pre-sexual stages in the cat intestinal epithelium (endopolygeny in merozoites). Endodyogeny is driven by assembly of two nascent cortical cytoskeletons in the cytoplasm of a mother cell (Anderson-White et al., 2012, Chen & Gubbels, 2013, Francia & Striepen, 2014, Nishi *et al.*, 2008, Goldman *et al.*, 1958). The cytoskeleton is composed of flattened vesicles known as alveoli organized in a quilt-pattern (Chen et al., 2016, Porchet & Torpier, 1977). On the cytoplasmic side the alveoli are supported by a meshwork of 10 nm filament forming proteins, which in turn are supported by a set of 22 cortical microtubules running in a spiral pattern from the apex to 2/3 the length of the parasite. This structure is also known as the Inner Membrane Complex, and many of its protein components are known as IMC proteins (Anderson-White *et al.*, 2011, Chen et al., 2015, Chen et al., 2016). The assembly of daughter parasite cytoskeletons nucleates at the centrosome outer core (Suvorova *et al.*, 2015) and progresses in an apical to basal direction throughout a 2-3 hr window in the 6.5 hr tachyzoite cell cycle (Nishi et al., 2008). The very apical end of the cytoskeleton is composed of a tubulin basket known as the conoid, which extrudes in a Ca^2+^-dependent fashion required for host cell invasion (Gonzalez Del Carmen *et al.*, 2009). A contractile ring at the basal end of the cytoskeleton known as the basal complex drives tapering of the nascent daughter cytoskeletons and in addition serves in maintaining a cytoplasmic bridge between divided parasites (Frenal *et al.*, 2017, Lorestani *et al.*, 2010).

One of the most intriguing structures in the *Toxoplasma* cytoskeleton comprises the apical annuli (aka peripheral annuli). The 5-6 annuli (meaning ring-shaped) were first observed with Centrin2 as ∼200 nm diameter rings residing at the transition between the single cap alveolus and the next more basal series of 5-6 alveolar vesicles (Hu *et al.*, 2006). Centrin2 is a small protein containing 4 EF hand domains that has been suggested to form filaments that contract in a Ca^2+^-dependent fashion (Hu, 2008). However, Centrin2 has multiple subcellular localizations: the centrosomes, the preconoidal ring, the apical annuli and the basal complex, and appears to fulfill several different functions in the parasite of which conoid localization was associated with microneme secretion (Lentini *et al.*, 2019). Recently, a protein was reported that only localizes to the annuli, Peripheral Annuli Protein 1 (PAP1), which is related to centrosomal proteins and harbors extensive coiled-coil regions (Suvorova et al., 2015). Since the abbreviation PAP is more commonly used for Phosphatidic Acid Phosphatases, we renamed PAP1 to Apical Annuli Protein 1 (AAP1).

To further our understanding of the apical annuli in *Toxoplasma* we resolved its proteome using proximity-based biotinylation proteomics (Roux *et al.*, 2012). A total of seven proteins were identified as *bona fide* apical annuli residents. Five of these, dubbed AAP1-5, shared low complexity and coiled-coil regions, suggestive of structural functions in the annuli. The AAP proteins are assembled into several concentric rings with diameters ranging from 200-400 nm. Centrin2 resides in the intermediate ring whereas an apical annuli methyltransferase (AAMT) is only present on the annuli in intracellular tachyzoites. Furthermore, the *aap* genes are conserved in the genomes of Coccidia that divide by internal budding but are absent from other Apicomplexa, which suggests that the annuli somehow facilitate internal budding. Knockout of AAP4, the most conserved AAP, demonstrated that the annuli are critical for *in vitro* expansion but not for the acute stage in mice. Overall, our data identify a set of novel proteins exclusively localizing to the apical annuli in *Toxoplasma,* which reveal the complex architecture and dynamics of this cytoskeleton assembly as well as several parallel insights toward putative function.

## Results

### 1. The annuli proteome mapped by proximity-based biotinylation

Proximity-based biotinylation is a powerful tool to analyze the protein composition of large complexes, in particular poorly soluble structures, such as the nuclear envelope (Roux et al., 2012), nuclear pore complex (Kim *et al.*, 2014), centrosome (Firat-Karalar *et al.*, 2014, Gupta *et al.*, 2015), and the cytoskeleton of *Toxoplasma* (Chen et al., 2015, Chen et al., 2016, Long *et al.*, 2017a, Long *et al.*, 2017b). We tagged Centrin2 with the small promiscuous biotin ligase BioID2 (Kim *et al.*, 2016) to create a merodiploid Ty-BioID2 fusion protein expressing parasite line. The fusion protein displayed the expected localization and the corresponding biotinylation in these structures upon addition of extracellular biotin (Fig. 1A, Fig. S1A). Mass spectrometry analysis of the purified biotinylated proteins identified several proteins in known Centrin2 localizations, including the basal complex (e.g. MyosinJ) and the centrosome (Centrin1, Centrin3) (Table S1). However, most hits were annotated as hypothetical proteins. To increase the identification of specifically biotinylated proteins, we expressed a morn1-driven BioID2-YFP construct in the cytoplasm of control parasites (Fig. S1B, C) and generated samples grown in absence of biotin for all used cell lines. We then calculated the normalized spectral abundance factor (NSAF) (Florens *et al.*, 2006) for all recovered proteins, to correct for overrepresentation of large proteins. By comparing rank-ordered proteins of the Centrin2 data set to the cytosolic control, we identified TGGT1_230340 with the greatest NSAF for a hypothetical protein in the Centrin2 data (Fig. S1D). We endogenously tagged TGGT1_230340 with a Myc-epitope tag and observed a signal towards the apical end of the parasite. TGGT1_230340 focused in 5-6 puncta around the apical cap, as judged by co-staining with β-tubulin (Fig. 1B). Co-localization with Ty-tagged Centrin2 confirmed the localization of TGGT1_230340 at the apical annuli (Fig. 1C).

**Figure 1.**
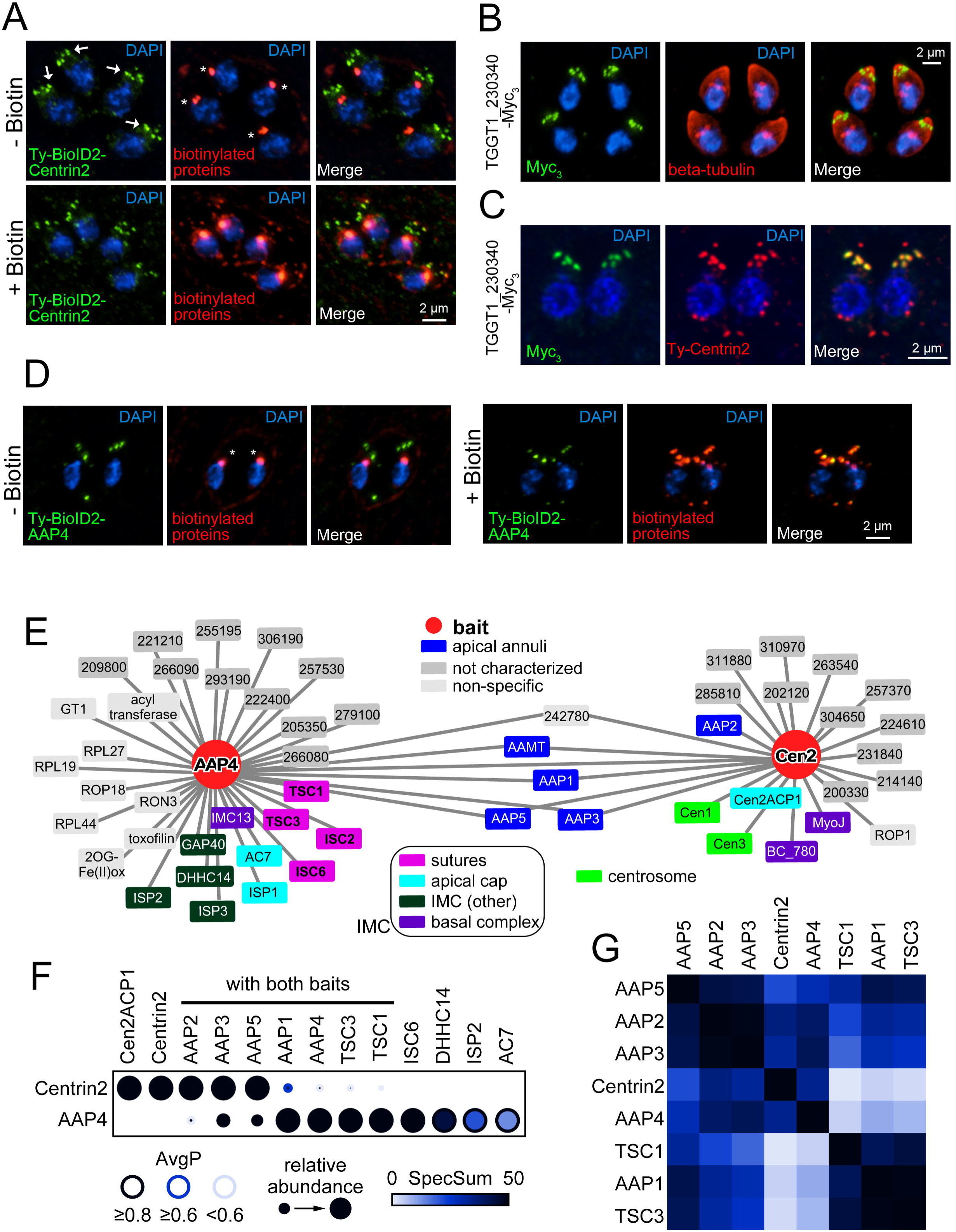
Protein-protein interaction (PPI) network analysis of the apical annuli. **A.** Episomal expressed Ty-BioID2-Centrin2 localizes predominantly to the apical annuli (arrows) but also to the preconoidal ring, the centrosome and the basal complex. Application of 150 μM biotin overnight facilitates biotinylation as detected with Streptavidin-594 (biotinylated proteins, red). Endogenous biotinylation of apicoplast proteins is always detected with Streptavidin (asterisks). **B.** Endogenously triple Myc-tagged (Myc_3_) TGGT1_230340 localizes towards the apical end of the parasites in 5-6 puncta. Parasites were co-stained with beta-tubulin (red) to highlight the parasite’s periphery. **C.** Airyscan imaging of TGGT1_230340 Myc_3_-tagged parasites transiently expressing Ty-Centrin2 (red). TGGT1_230340 only co-localizes with Centrin2 at the apical annuli and is not present in other subcellular Centrin2 localizations. Due to its localization we named TGGT1_230340 apical annuli protein 4 (AAP4). **D.** AAP4 was endogenously tagged with Ty-BioID2 at the N-terminus (see Fig. S1E). Upon application of 150 μM biotin, increased biotinylation of the apical annuli can be detected by Streptavidin staining (biotinylated proteins, red). Endogenously biotinylated proteins of the apicoplast are always detected (asterisks). Blue: DAPI stain. **E.** PPI networks were modeled by calculating probabilistic bait-prey interactions using SAINTexpress (Lambert et al., 2015) and plotted with Cytoscape (Saito et al., 2012, Shannon et al., 2003) in a ball-and-stick model. Preys with an AvgP (average individual probability for SAINTexpress analysis) ≥ 0.5 are shown for each bait. **F.** The statistical support for the core protein set of the apical annuli is shown as a dot plot generated with ProHits-viz (Knight et al., 2017). Note that the relative abundance compares between the two samples and not within an individual sample (e.g. AAP4 was one of the most abundant proteins identified in Centrin2 BioID data, but compared to the number of spectra in the AAP4 data set it deceivingly appears to be relatively low abundant in the Centrin2 data set). Also note that AAP2 is present in the AAP4 BioID data set and TSC1 and TSC3 in the Centrin2 BioID data set, but only with low AvgP scores (< 0.2). AvgP: average individual probability for SAINTexpress analysis, SpecSum: sum of all spectra for an individual protein. **G.** The data set was further assembled into a prey-prey correlation map showing the apical annuli core set. The distance between individual preys is expressed by color, Black indicates correlating preys, non-correlating preys are shown in white. See Table S1 for the complete data.

To gain further insights into the enigmatic apical annuli we reciprocally tagged this new apical annuli protein, dubbed AAP4, with BioID2 on the N-terminus. To achieve this we utilized the selection-linked integration (SLI) strategy previously developed in *Plasmodium falciparum* (Birnbaum *et al.*, 2017). We inserted the HXGPRT selectable marker linked to Ty-BioID2 by a T2A skip-peptide at the 5’end of the endogenous *aap4* ORF (Fig. S1E). This line showed the previously observed subcellular localization for AAP4 and showed increased biotinylation upon BioID activation (Fig. 1D, Fig. S1F).

Upon mass spectrometry and further analysis with SAINTexpress software (Teo *et al.*, 2014), nearly 1/3 of the proteins in the AAP4 BioID2 data set were known to localize to various aspects of the IMC. Notably transversal suture components (TSC) 1 and 3 and IMC suture component (ISC) 2 and 6 (Chen et al., 2015, Chen et al., 2016) stand out (Fig. 1E). However, the largest number of the proteins specific to AAP4 were annotated as hypothetical (Table S1).

We used the mass spectrometry data of both Centrin2 and AAP4 BioID experiments to assemble a protein-protein interaction (PPI) network. Subsequently this data set was analyzed for probabilistic bait-prey interactions that have an AvgP (average individual probability for SAINT analysis) ≥ 0.5 (Fig. 1E) (Choi *et al.*, 2011, Lambert *et al.*, 2015). Several proteins are shared between the Centrin2 and AAP4 data sets, of which the following localized exclusively to the apical annuli: AAP1 (TGGT1_242790) previously identified in (Suvorova et al., 2015); AAP3 (TGGT1_313480); and AAP5 (TGGT1_319900) (Fig 2A). A fourth shared protein is annotated as a putative methyltransferase (TGGT1_310070) that we named apical annuli methyltransferase (AAMT). A fifth, hypothetical protein (TGGT1_242780) shared between Centrin2 and AAP4 is reminiscent of a translation-related factor and we considered it a non-specific hit. To statistically support our observations we visualized the analysis with a dot plot that highlighted the SAINTexpress readout. This dot plot is composed out of the AvgP, the relative protein abundance of a given protein seen in the respective BioID experiment and the sum of detected spectral counts (SpecSum) of that protein obtained in all individual replicates (Fig 1F).

**Figure 2.**
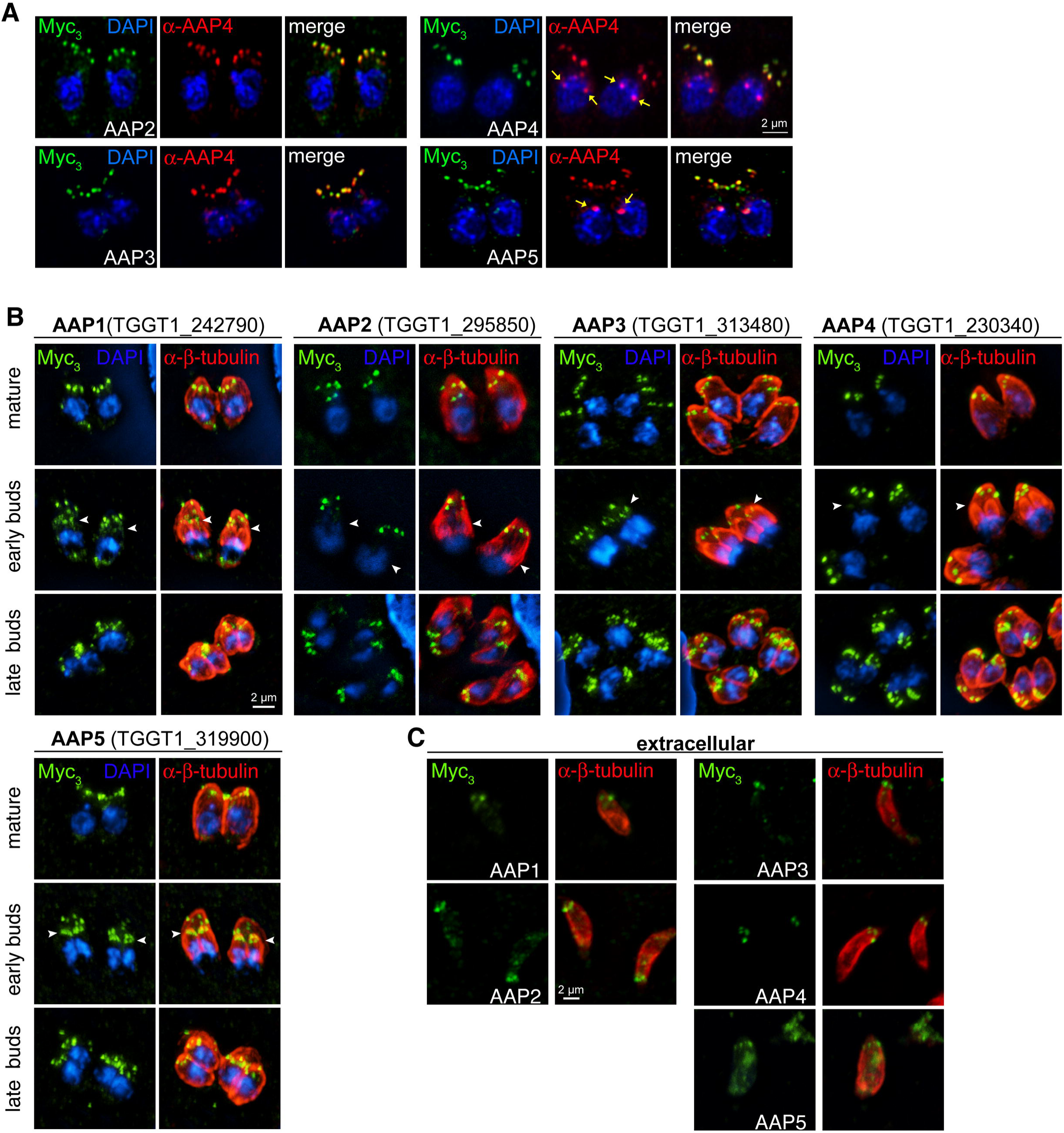
Localization and dynamics of the apical annuli proteins (AAPs). **A.** AAP genes were endogenously tagged with a triple Myc-epitope tag (Myc_3_) and co-localized with a specific antisera that recognizes AAP4. Images were acquired on the Airyscan system. Yellow arrows indicate cross-reactive signal seen with the AAP4 antiserum close to the nucleus (see also Fig 6A and Fig S5B). **B.** Protein dynamics of identified AAPs were further followed along the intracellular division cycle of the tachyzoite. The periphery of the parasite is visualized with Tg-β-tubulin antiserum staining (red). Arrowheads in the middle plane indicate annuli presence in early daughter buds. All AAPs are present in late daughter buds when the mother’s cytoskeleton is being degraded. Blue: DAPI stain. **C.** AAPs localize to the apical annuli in extracellular parasites, although AAP5 only exhibits a weak signal.

This highlighted an additional protein shared between the two data sets, AAP2 (TGGT1_295850), which upon validation also localizes to the annuli (Fig. 2A). However, the statistical support in the AAP4 sample is very low, in contrast to its robust signal in the Centrin2 data. Furthermore in this dot plot, AAP4 appears to be relatively poorly represented in the Centrin2 data set. This representation is skewed by the extremely high peptide counts of AAP4 in the AAP4 sample itself, resulting in a relatively low abundance in the Centrin2 sample in this comparative representation. Absolute numbers put AAP4 in the top 3 of overall identified proteins in the Centrin2 sample. Reciprocally, the AAP4 sample revealed proteins found in the IMC sutures that are of relatively low abundance in the Centrin2 sample. This indicated that the annuli might be embedded in the IMC sutures. We further used the data to assemble a prey-prey correlation heat map of the interactions (Fig. 1G). This identified several clusters of annuli and suture proteins that overlapped with each other. For example, identical behavior is suggested for AAP2 and AAP3, which could indicate a coupled function. This further closely resembles the AAP5 pattern, suggesting a close proximity of these three proteins in the parasite. Likewise, AAP1, TSC1, and TSC3 appear to be closely associated, whereas AAP1 connects to the AAP2/3/5 complex. Centrin2 and AAP4 appear to sit slightly apart (likely due to their skewed abundance in the data set as they were used as baits) but seem to be closer to the AAP2/3 complex than to the AAP1/TSC1/3 complex. Thus, these data provide a tentative architectural layout of the annuli and how they interface with the IMC sutures.

### 2. Experimental validation of AAP proteins predicted by PPI analysis

We validated the BioID2-analysis based AAP annotation for the AAP2-5 genes by tagging the 3’-end of the endogenous loci with a triple Myc-epitope tag (Myc_3_) and immunofluorescence assays (Fig. 2, Fig. S2A, B). We observed apical annuli localization for all AAP proteins as judged by co-staining with an AAP4 antiserum that we generated (Fig. 2A). In addition we tagged the previously identified AAP1 (Suvorova et al., 2015) with a triple Myc epitope tag using the SLI system at the 3’end (Fig. S2C). We observed differences in signal intensity between mother and daughter buds for the different AAPs (Fig. 2B). By relative intensity to the mother, AAP5 is most enriched in the daughters followed in sequence of diminishing intensity by AAP1, AAP3, AAP4 and AAP2. These data correspond to observations of Centrin2, which was reported to be present in the annuli when the daughter cytoskeleton is being assembled (Hu et al., 2006). Our data expand on this observation by suggesting a putative hierarchical protein assembly of the annuli. We did not observe comparable changes in annuli association of AAP1-4 in extracellular parasites, except for AAP5, for which the signal sharply reduced in intensity (Fig. 2C).

### 3. The AAPs are pluriform coiled-coil domain containing proteins

We queried ToxoDB and performed other database searches to reveal functional features in the primary sequences of the AAPs. The AAPs are of a variety of size, domain composition and phosphorylation status in the tachyzoite phosphoproteome (Treeck *et al.*, 2011) (Fig. 3A-C). We identified coiled-coil regions as previously reported for AAP1 (Suvorova et al., 2015) in all of them except AAP5. AAP5 contained no recognizable domain features except an α-helix-rich region. The presence of coiled regions however, appears to be the most shared feature across the AAPs. AAP1 was first described as centrosomal CEP250-related protein of approximately 200 kDa protein (Suvorova et al., 2015). AAP1 stands out from the other AAPs in that no phosphorylation sites were detected. A <u>H</u>istidine kinases, <u>A</u>denyl cyclases, <u>M</u>ethyl-accepting proteins and <u>P</u>hosphatases (HAMP) linker domain is present in the center of AAP1 (Fig. 3B). In bacteria, HAMP linkers are transmembrane two-component sensors that form dimers composed of four-helical bundles and can exist in two conformations to transduce signals (Bhate *et al.*, 2015). In fungi, HAMP domains are critical in histidine kinases of fungicide targets as well as have been shown to function in osmo-sensing (Defosse *et al.*, 2015, Meena *et al.*, 2010). However, the *Toxoplasma* kinome neither contains any predicted histidine kinase (Peixoto *et al.*, 2010) nor does AAP1 contain a transmembrane domain, thus suggesting a function distinct from the well-characterized, HAMP domain containing systems. Furthermore, detailed analysis of AAP1’s coiled-coil region identified 11 repeats of a 33 amino acid sequence (Fig. 3B, D). Stretches of K and E residues make this a highly charged repeat. The repeats are predicted to form α-helical coiled-coils interspersed by a short linker region (Fig. 3B). Similar highly charged repeats are found in three axoneme-associated protein mst101(1-3) of *Drosophila hydei* (Neesen *et al.*, 1999, Neesen *et al.*, 1994). The 16 amino acid repeats in DhMst101 also contain regularly spaced cysteine-residues that are expected to form long alpha-helical rods cross-linked by numerous Cys-Cys bridges. Dhmst101 proteins are part of the outer sheath of the sperm tail where they presumably help to provide a tight but elastic envelope for the extremely extended (20 mm) spermatozoa of *D. hydei*. However, Cys residues are not present in AAP1 suggesting the putative rods formed by this repeat are not cross-linked.

**Figure 3.**
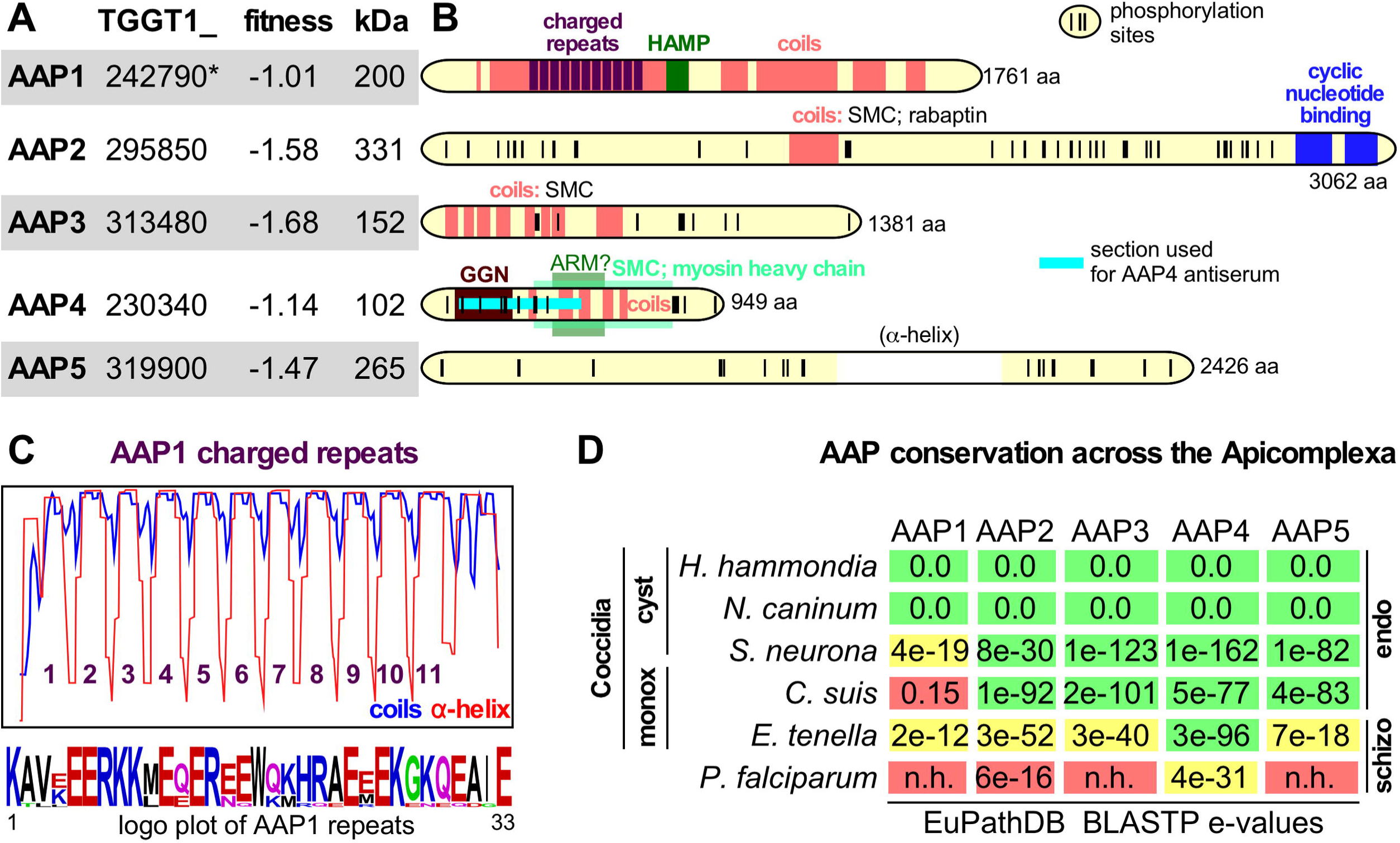
Annotation and sequence analysis of the AAPs. **A** AAP annotation and functional data available through ToxoDB.org (Gajria *et al.*, 2008). Fitness scores defined by genome-wide CRISPR screen for fitness across three lytic cycles (Sidik et al., 2016). *In GT1 annotation AAP1 comprises two genes: TGGT1_242790A (fitness score 0.02, with an extended N-terminus) and TGGT1_242790B (fitness score −1.01). **B.** Domain annotations of AAP1-5 made through searches on ToxoDB, SMART, and PFam databases, NCBI Nr PBLAST searches and coiled-coil predictions (coils window size of 28; 100% probability predictions are shown). The long, largely α-helical domain in AAP5 is between brackets as this feature is not displayed for AAP1-4, but it is the only distinguishable feature in AAP5. Phosphorylation sites detected in the tachyzoite phosphoproteome are marked with vertical ticks (Treeck et al., 2011). The number at the C-terminus indicates the number of amino acids. **C.** Coils prediction by the coils server using a window size of 21 overlaid with the PSIPRED predicted α-helical repeats (ToxoDB) identify 11 repeated regions (y-axis represents 0-100% probability range of α-helix or coiled-coil), whose consensus repeat sequence is provided in the logo plot (Crooks *et al.*, 2004) at the bottom (see also Fig S2). **D.** Conservation of the AAP proteins throughout the Apicomplexa through BLASTP of the *Toxoplasma* AAPs against EuPathDB (Aurrecoechea et al., 2013) and OrthoMCL (Chen et al., 2006). Colors represent likelihood of functional conservation based on manual assessment of the quality and length of sequence alignments and genomic synteny (green: robust ortholog; yellow: putative ortholog; red: no ortholog). cyst: cyst forming; monox: monoxenic n.h.: no homology; endo: asexual division by endodyogeny and/or endopolygeny; schizo; asexual division by schizogony. The absence of tissue cysts in *C. suis* is likely a secondary loss in this lineage as it is phylogenetically is more closely related to *Toxoplasma* than to *Sarcocystis* (Carreno *et al.*, 1998).

Using the annotated apicomplexan genomes assembled on EuPathDB (Aurrecoechea *et al.*, 2013) and OrthoMCL (Chen *et al.*, 2006) databases we determined the AAP gene conservation across the Apicomplexa. Gene orthologous to AAP1-5 were easily found in the closest relatives of *Toxoplasma, Hammondia hammondi* and *Neospora caninum* (Fig 3D). The AAP1 orthologs in *Hammondia hammondi* and *Neospora caninum* both contain very synonymous charged repeats though the number of repeats is 10 in NcAAP1 and 8 in HhAAP1 (Fig. 3D, Fig. S3). AAP1 is poorly conserved beyond these close relatives as a robust ortholog is not detectable in the next closest relatives *Sarcocystis neurona* and *Cystoisospora suis*.

AAP2 is the largest of the AAP proteins with a predicted mass of 331 kDa. Sequence analysis revealed two C-terminal cyclic nucleotide (cNMP) binding domains (Fig. 3A). In addition, the center of the protein harbors a coiled-coil domain, which presented weak homology to a variety of coiled-coil domains found in Structural Maintenance of Chromosomes (SMC) proteins as well as to a protein known as Rabaptin. Rabaptin is a coiled-coil protein that interacts with Rab5 and functions in endosomal vesicle recycling by facilitating membrane docking and fusion (Stenmark *et al.*, 1995, Deneka *et al.*, 2003). Seventy-three phosphorylated residues were detected in the AAP2 phosphoproteome, which aggregate in 2-3 clusters outside the recognizable functional domains (Fig. 3A). Across the Apicomplexa AAP2 stands out with homology found in *E. tenella* (1819 aa) sharing the coiled helix domain and the cNMP binding domains, but this protein is otherwise not homologous, not syntenically organized in the genome and might have evolved fast potentially toward a different function. The protein with the highest AAP2 e-value BLASTP in the *S. neurona* is nearly twice the size (3658 aa) and does not contain the cNMP binding domains, which could be due to a misannotation as it is syntenically organized in the genome (Fig. 3D).

AAP3 is predicted to be 152 kDa and harbors a series of coiled-coil domains, which displayed weak and likely random homology to SMC proteins (Fig. 3A). In addition, 15 phosphorylated residues were detected in the tachyzoite phosphoproteome (Fig. 3B). One clear ortholog is present in *S. neurona* and *C. suis,* but not present in *E. tenella* (Fig 3D).

AAP4 is the smallest AAP protein with a predicted molecular weight of 102 kDa containing 26 detected phosphorylation sites, seven of which are clustered in the C-terminus whereas the others are located in the N-terminal half (Fig. 3A, B). A series of coiled-coil domains make up the central parts of the protein and display some tentative homology to an Armadillo-like domain (ARM) as well as the coiled-coil tail of myosin heavy chains, suggestive of a structural function. Furthermore, toward the N-terminus a domain with homology to the GGN superfamily is present. GGNs or gametogenetins, are present in mammalian sperm cells, membrane associated, and have likely function in vesicular trafficking toward sperm maturation (Lu & Bishop, 2003, Zhao *et al.*, 2005). Thus, the domains present in AAP4 provide a hint toward a potential role in membrane trafficking for the apical annuli. AAP4 is by far the strongest conserved AAP protein across the Apicomplexa as a putative ortholog is even detected in *Plasmodium falciparum* (Fig. 3D).

The predicted AAP5 protein size is 265 kDa in which no specific domain features could be detected at all except a centrally located section almost exclusively composed of an α-helix. The tachyzoite phosphoproteome reported 36 phosphorylation sites on AAP5.

### 4. A methyltransferase resides on the apical annuli in intracellular tachyzoites

To validate the localization of the methyltransferase AAMT, identified by our PPI network in Figure 1 E-G, we tagged the endogenous locus with a triple Myc-epitope tag (Myc_3_) (Fig. 4A, Fig. S2A, B). We did observe a signal in the apical annuli, but found AAMT also at the apical end of the parasite and present in forming daughter buds (Fig. 4A, arrowheads). The apical annuli localization was confirmed by co-localization with AAP4 antiserum, but is not as complete as observed for the AAPs (Fig. 4B). Furthermore, in extracellular parasites the apical annuli localization changed and we observed a more diffuse localization somewhat reminiscent of the IMC sutures, in particular the transverse sutures (Fig. 4C). These data indicate that the apical annuli could be a signaling platform associated with the transition between the intracellular and extracellular states of the tachyzoite.

**Figure 4.**
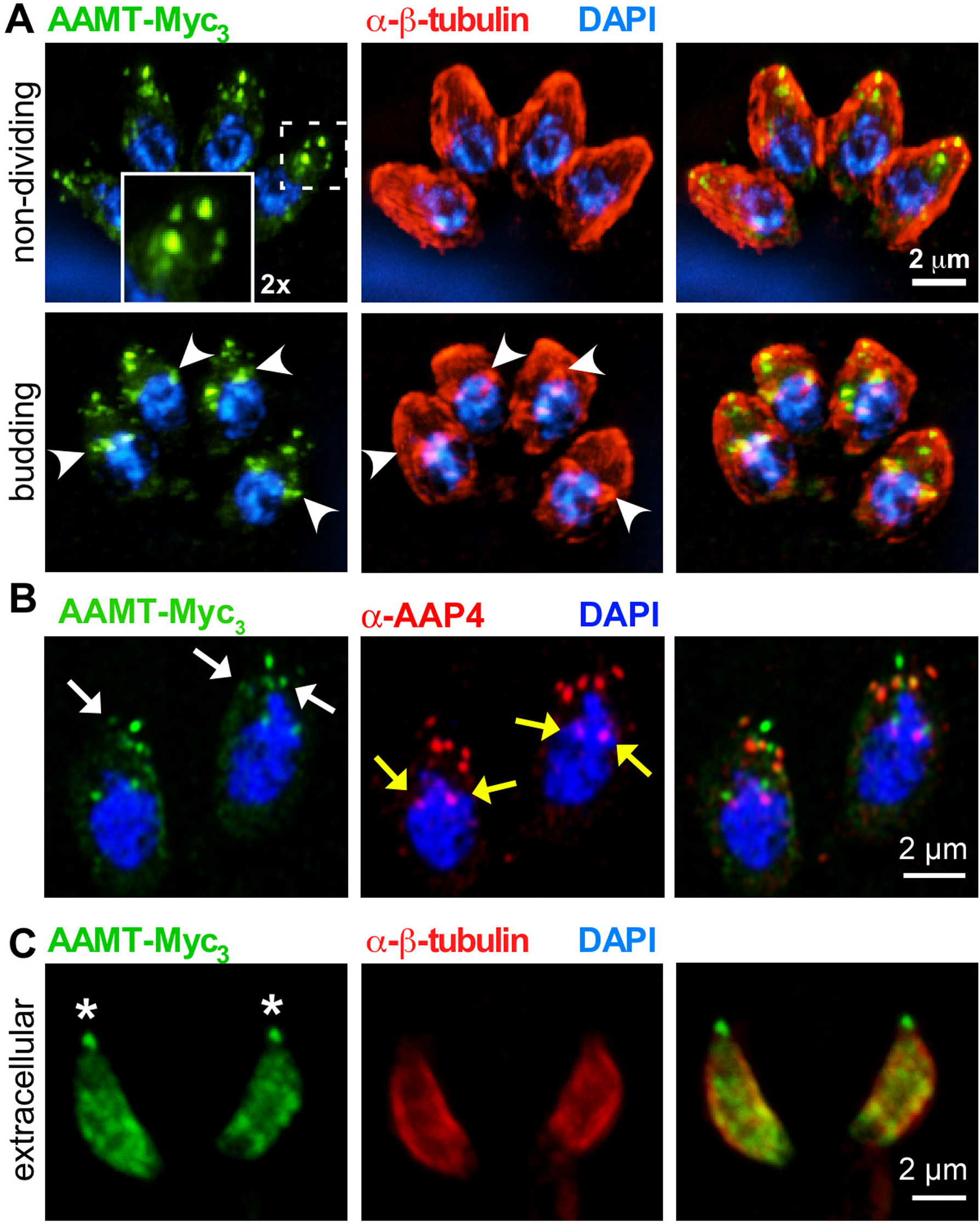
The methyltransferase AAMT localizes to the annuli in intracellular parasites. **A.** Endogenously triple-Myc tagged (Myc_3_) AAMT displays apical annuli localization in mature parasites (top panels, boxed area magnified 2-fold in inset). We further detected AAMT signal association with the conoid and in forming daughter buds (lower panels, marked by arrowheads).. **B.** Airyscan imaging of AAMT-Myc_3_ parasites co-localized with AAP4 antiserum (red). White arrows indicate the apical annuli, as highlighted by AAMT and AAP4 staining. Note that the AAMT localization to the annuli is not as complete as detected for other AAPs. Yellow arrows indicate cross-reactive signal seen with the AAP4 antiserum close to the nucleus (see also Fig 6A and Fig S5B). **C.** In extracellular parasites AAMT re-distributes to the (transverse) IMC sutures but remains associated with the conoid (asterisks). Tg-β-tubulin serum (red) (Morrissette & Sibley, 2002) highlights the cortical cytoskeleton. Blue: DAPI stain.

### 5. The apical annuli components form differently sized concentric rings

Immuno-electron microscopy of Centrin2 revealed that the signals in individual annuli were 200 nm diameter rings (i.e. annuli) (Hu et al., 2006). By conventional wide field microscopy we observed some of the AAP signals as ring-like whereas others appear as solid dots suggestive of sub-domains in the annuli (Fig. 2B). To further resolve the architecture of the annuli we applied super-resolution structural illumination microscopy (SR-SIM). As seen in Figure 5A and Movie S1, 2, we observed signals of different size and shape for the AAPs and Centrin2. Measuring the diameters of individual annuli lead to the resolution of three groups with statistically different diameters (Fig. 5B). The widest diameter of 394 nm was observed for AAP4, followed by 305 nm for the combination of AAP5 and Centrin2, 255 nm for AAP3 and 202 nm for AAP2. With this SR-SIM approach we resolved clearly visible ring-like signals for AAP4 and Centrin2. The AAP4 signal appears to have a squarish shape with accumulation on four corners. However, we cannot exclude potential distortion caused by the SR-SIM image processing or parasite fixation and are cautious to draw a firm conclusion based on these images. Ring-like structures were inconsistent for AAP3 and AAP2, but we found examples even for the smallest of the AAPs, AAP2 (Fig. 5C). Moreover, AAP2-Myc_3_ co-localization with AAP4 antiserum displayed that AAP2 indeed resides within the AAP4 ring (Fig. 5C). To further characterize AAPs in the parasite, we co-localized AAP4 with ISC2, a previously described component of the IMC sutures (Chen et al., 2015). This co-localization showed AAP4 signal apical to ISC2, where the IMC borders the apical cap (Fig. 5D, Movie S3). Combing our BioID data with the SR-SIM results, we resolved the annuli as a multilayered assembly that consists of several AAPs of different diameter. The specific enrichment of structural components of the IMC sutures, seen by BioID (e.g. TSC1 and 3, ISC2 and 6, ISP1 and 2) shows the integration of the annuli between the IMC plates at the apical end of the parasite (Fig. 5 E).

**Figure 5.**
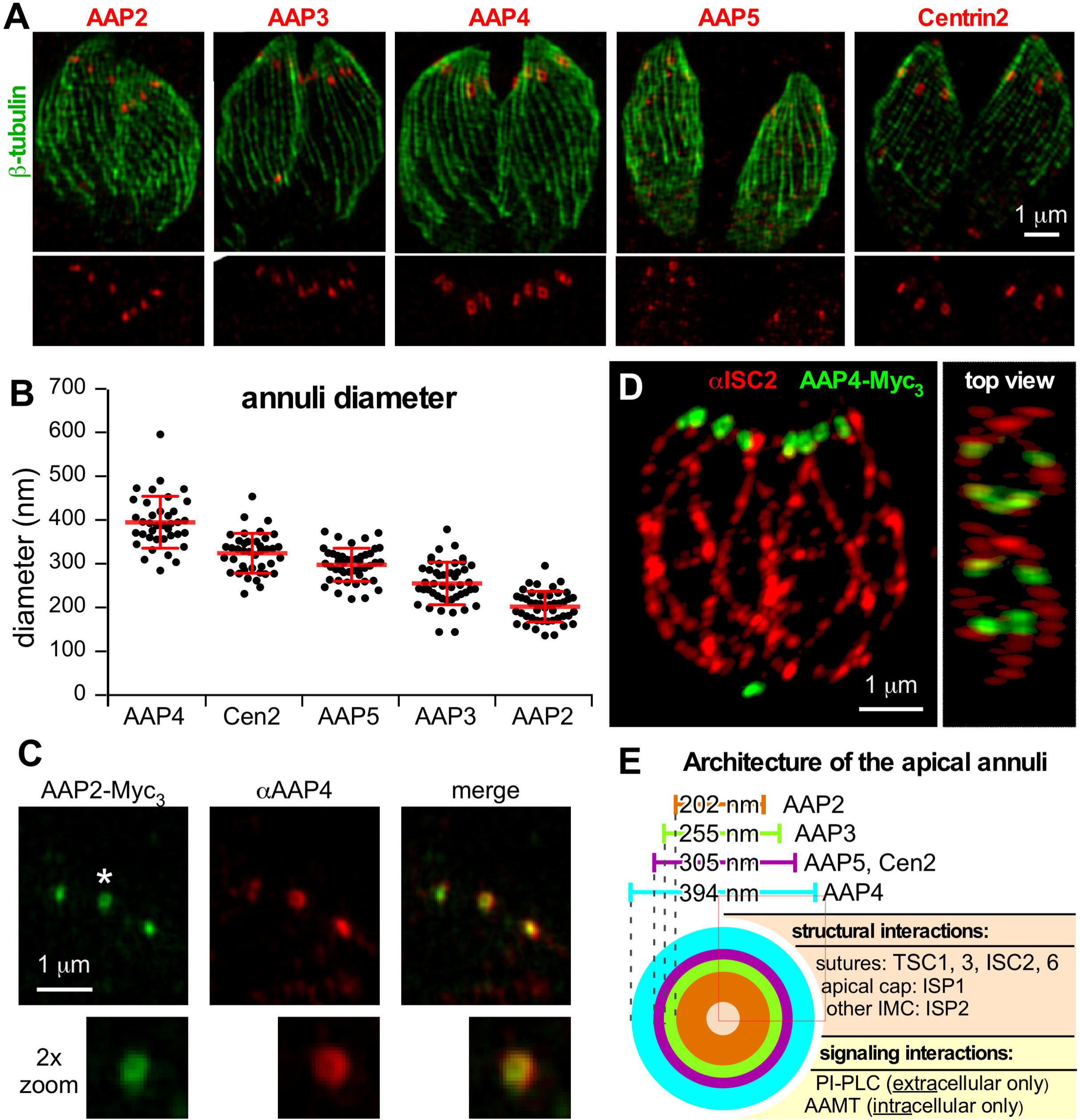
The apical annuli display a concentric ring architecture. **A.** SR-SIM analysis of endogenously Myc-tagged AAP and Centrin2 parasite lines co-stained with Tg-β-tubulin antiserum (Morrissette & Sibley, 2002). The lower 50% of the imaged stack was combined into a Z projection for each image. See Supplementary Movies S1 and S2 for a 3D-reconstruction of the entire AAP4 image stack. **B.** Quantification of AAP donuts. Diameters of at least 40 individual annuli were measured for each cell line. Error bars represent standard deviation. Statistics: paired two-tailed *t*-test analysis indicated that all signals except AAP5 and Centrin2 (*p*=0.485) are significantly different from each other (*p*<0.0001). **C.** SR-SIM analysis of C-terminally triple-Myc tagged AAP2 co-stained with a specific antiserum recognizing AAP4. AAP2 signals, shown to exhibit the smallest average annuli diameter, were observed as donuts and are embedded by the AAP4 signal. Image is cropped to the apical section of a tachyzoite. **D.** SR-SIM analysis of C-terminally triple-Myc tagged AAP4 co-stained with a specific ISC2 antiserum. The annuli, highlighted by AAP4, localize to the apical end of the IMC sutures (ISC2 signal). See Supplementary Movie S3 for a 3D-reconstruction of the entire image stack. **E.** Schematic presentation of the apical annuli architecture incorporating microscopy and PPI data. PI-PLC interaction is gleaned from (Hortua Triana et al., 2018).

### 6. The apical annuli contribute to pathogenesis

The fitness scores of the AAP genes in the genome-wide CRISPR screen are between −1.01 and −1.68, which is suggestive of non-essential functions during the lytic cycle of *Toxoplasma* (Sidik *et al.*, 2016) (Fig. 3A). However, to directly test the biology of the annuli we depleted AAP2 and AAP4 as they represent the smallest and largest ring-like signals, respectively. Furthermore, AAP4 is of particular interest as it harbors the most functional domains and is the most conserved among the Apicomplexa. We established a tetracycline-regulatable promoter replacement for AAP2 (cKD), which did not result in any loss of tachyzoite fitness as assessed by plaque assays (Fig. S4D). For AAP4 we established a complete gene knockout parasite line (Fig. S5A) and monitored loss of AAP4 using the AAP4 antiserum by IFA and in western blots (Fig. 6A, Fig. S5B). The AAP4-KO line proliferated significantly slower than the parental line (Fig. 6B-C), and this proliferation defect was restored upon genetic complementation with the *aap4* gene, which excludes potential artifacts of genome editing (Fig. 6, Fig. S5). To test the role of annuli *in vivo*, we infected C57BL/6 mice with AAP4-KO, parental RHΔKu80 (control), and AAP4-KO complemented parasites. As expected, mice infected with either the control line or the AAP4-KO complementation succumbed to the infection within 8 days (Fig. 6C). AAP4-KO infected mice survived acute infection one day longer, until day 9 (Fig. 6C). In a replication experiment using 100 parasites for infection we observed a similar trend for mice infected with AAP4-KO parasites, surviving infection one day longer. However, the weight change pattern was comparable to mice infected with control and AAP4-KO complemented parasites (Fig. S6). Together these results suggest that the apical annuli have only a minimal role in acute virulence of *Toxoplasma*

**Figure 6.**
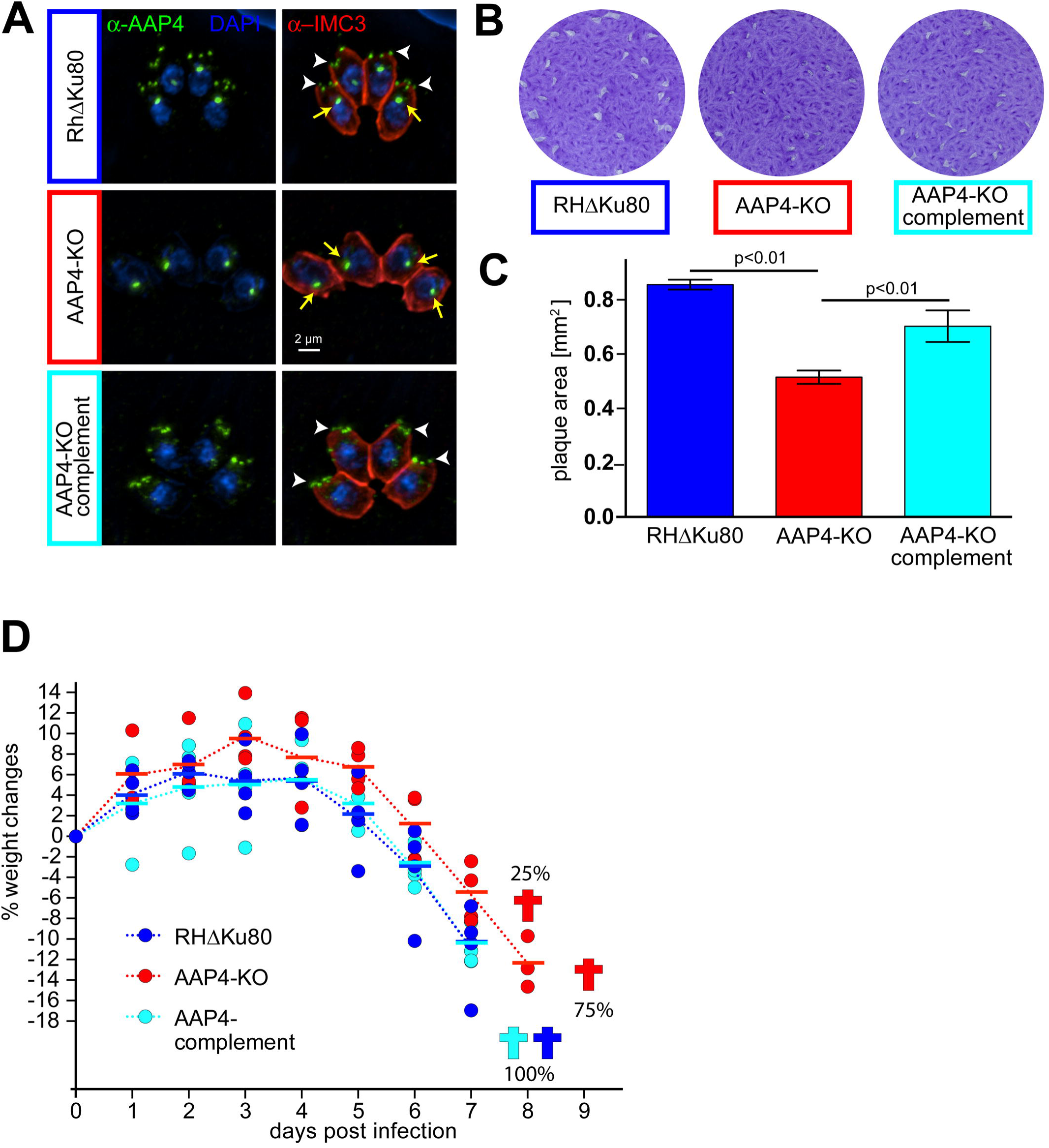
Knockout analysis of AAP4 reveals decreased fitness *in vitro* and reduced virulence *in vivo*. **A**. IFA using specific α-AAP4 serum (green). White arrowheads indicate AAP4 signal at the annuli, yellow arrows indicate a cross-reactive signal seen close to the nucleus (see also Fig S5B). Parasite periphery is shown by α-IMC3 serum (red). Blue: DAPI stain. **B.** Representative plaque assays of AAP4-KO parasites after seven days of growth. AAP4-KO parasites form significantly smaller plaques compared to control (RHΔKu80) or AAP4-KO complement parasites. **C**. Quantification of plaque assays. Three biological replicates are shown; 40-80 independent plaques per condition and experiment were quantified; error bars represent standard deviation; For statistical analysis an one-way analysis of variance (ANOVA) test was applied and significance determined with a post-hoc Tukey’s honestly significant difference (HSD) test. **D.** C57BL/J6 mice infected with 1000 AAP4-KO parasites survive one day longer than mice infected with equal numbers of control (RHΔKu80) or AAP4-KO complemented parasites. Weight changes relative to the starting day (day 0) are shown for each of the four mice per group (round symbols). Horizontal bars represent the group average. Weight change patterns did not show significant differences between the groups, as tested by one-way ANOVA test. An additional infection experiment with a 100-parasite inoculum is present in Fig. S6.

## Discussion

Our interrogation of the apical annuli composition by a proximity-based biotinylation approach with subsequent statistical analysis using Centrin2 and AAP4 as baits, identified seven proteins localizing to this structure. We identify four new annuli proteins, AAP2, AAP3, AAP5, and the AAMT. AAP4 has been previously detected in BioID data sets of the alveolar suture component ISC4 (Chen et al., 2016), of the conoid (Long et al., 2017a) and a protein above the conoid, SAS6L (Lentini et al., 2019). Lentini et al. further co-localized AAP4 with Centrin2 and describe a relationship between Centrin2 knockdown and AAP4 abundance in the parasite (Lentini et al., 2019).

Recently, a minor population of TgPI-PLC was also reported to reside on the apical annuli (Hortua Triana *et al.*, 2018). We did not detect TgPI-PLC in our BioID experiments, which is presumably due to its dynamics. TgPI-PLC only localizes to the annuli in extracellular parasites. Interestingly, we identified AAMT as an apical annuli component only in intracellular parasites, thus exhibiting the exact opposite dynamics of TgPI-PLC. Furthermore, AAMT is widely conserved across the Apicomplexa, but its relative fitness score of −1.22 suggests it likely does not fulfill an essential function (Sidik et al., 2016). This is in contrast to the essential apical complex lysine methyltransferase AKMT (Heaslip *et al.*, 2011), which localizes to the conoid and is released into the cytoplasm to activate gliding motility (Jacot *et al.*, 2016). Further work is required to assess if AAMT participates to this process, as the localization dynamics of both methyltransferases are similar. The genome-wide CRISPR/Cas9 screen throughout three lytic cycles further indicates that none of the AAPs significantly affects parasite fitness (Sidik et al., 2016). We indeed observed no fitness defect in AAP2-depleted parasites. And although parasites devoid of AAP4 have decreased fitness *in vitro*, acute virulence is only minimal affected as indicated by our *in vivo* experiments. Further work is needed to assess whether the various AAPs have redundant functions in maintaining annuli structure and/or function.

Our data define the previously unknown architecture of the apical annuli from three different perspectives: 1. the concentric rings observed by super-resolution microscopy; 2. the BioID approach, that identified several alveolar suture proteins; 3. the domains contained in the AAP proteins. Previous immunoelectron microscopy on Centrin2 reported a diameter of about 200 nm for individual annuli (Hu et al., 2006). By SR-SIM we came to a Centrin2 diameter of about 300 nm, a difference that could be attributed to variations in the fixation method and/or the size and placement of eGFP (Hu et al., 2006) vs. Myc-epitope tag (this study). We consistently used the same tag and same imaging method on intracellular parasites; therefore, our data is consistent along all analyzed AAPs in this regard. AAP2 forms the smallest structure (∼200 nm) with AAP3 (∼250 nm) and AAP5/Cen2 (300 nm) in between and AAP4 at the upper end of the scale (∼400 nm) (Fig 5E). The smaller AAP2 and AAP3 ring-like openings were not consistently observed. This may be a result of the angle the annuli were imaged at or because their diameter being near the resolution limit. However, we cannot exclude the alternative explanation that the annuli opening might be dynamic depending on the conditions, with Centrin2 providing a robust candidate for the contractile force (Hu, 2008). Obtaining consistent images of AAP1 on the annuli was challenging, suggesting AAP1 might not be tightly associated with the annuli. The putative rod-forming domain repeats in AAP1 likely serve a structural function, but given the relatively weak or peripheral presence in the annuli suggests it might not be critical for the annuli.

Secondly, the BioID data contained several significant IMC localizing proteins like ISP1 and ISP2, which are palmitoyl-anchored in the cap alveolus and central alveoli, respectively (Beck *et al.*, 2010). However, most abundant were proteins found in the sutures between the alveolar plates, which is consistent with the position of the annuli at the edge of the cap alveolus. Along this line, the first report of TSC1 (named SIP then) indicated a reasonable level of Centrin2 co-localization at the apical annuli, besides other suture localizations (Lentini *et al.*, 2015). The localization of AAP4 at the apical edge of the lateral ISC2 suture signal further pinpoints that the annuli sit right at the intersection where two lateral alveoli meet the cap alveolus. These data suggest that the annuli are actually embedded in the sutures, most likely at the intersections between the cap alveolus and where the 5-6 central alveolar plates meet each other.

A suture-embedded annuli model fits with absence of transmembrane domains or other membrane anchors on the AAP proteins. Our structural analysis highlights the coiled-coil domains as the key feature in AAPs, with the exception of AAP5. These domains are typically present in fast evolving proteins (Kuhn *et al.*, 2014) and also found in centrosomal proteins (e.g. AAP1 was initially identified for its centrosomal protein features (Suvorova et al., 2015)). In this regard, it is of note that desmosomes in suprabasal epidermal cells associate with centrosomal proteins and CLIP170 to cortically organize microtubules (Sumigray & Lechler, 2011). Desmosomes are present in multicellular organisms and mediate tight interactions between epithelial cells of tissues exposed to mechanical stress (Garrod & Chidgey, 2008, Thomason *et al.*, 2010, Nekrasova & Green, 2013). Interestingly, the appearance and composition of the apical annuli is somewhat reminiscent of desmosome features. Of course, the presence of Centrin2 further supports the close relationship between the annuli and centrosome. This parallel between desmosomes and apical annuli suggests a putative structural function for the annuli, as they are in apposition of both the IMC and subpellicular microtubules. However, the number of 5-6 annuli does not appear to be geometrically related to the 22 sub-pellicular microtubules and as discussed above fits better with the 5-6 central alveolar plates.

The AAPs are a set of diverse proteins harboring coiled-coil domains with hints of both structural and signaling functions putatively regulated by extensive phosphorylation. Another glimpse toward their potential function comes from their conservation across the apicomplexan lineage (Fig 3D). They are narrowly conserved in a subset of the Coccidia that form cysts (*Hammondia, Neospora*, and *Sarcocystis* spp.). Interestingly, our conservation analysis also identifies AAP homologs in the non-cyst forming, monoxenous *Cystoispora suis*, however, the absence of cyst-forming capacity is a secondary loss of this feature (Worliczek *et al.*, 2013). The consistently shared phenomenon in this sub-group of Coccidia is that they multiply by an internal budding process (endogeny). This in contrast to most monoxenous Coccidia like the *Eimeria* spp., as well as most other Apicomplexa infecting humans or livestock, which multiply by a cortical budding process dubbed schizogony. A key difference in context of annuli function is that the cortical cytoskeleton is absent from the mother and is maintained during endogeny, whereas this is disassembled shortly after host cell invasion in parasites dividing by schizogony (Anderson-White et al., 2011, Dubey *et al.*, 2017). We hypothesize that this mother IMC during trophozoite expansion poses an obstacle for exchange of nutrients and waste products across the plasma membrane with the extracellular (vacuolar) environment. Thus, the apical annuli could presumably serve a pore function facilitating efficient exchange across this barrier.

In addition to their putative structural and/or pore-forming functions, the annuli might be a platform for signaling events. Support is found in the differential localization of TgPI-PLC and AAMT in intracellular and extracellular parasites, and in the presence of protein domains with a signaling signature found in some of the AAPs. Of note are the HAMP domain in AAP1, which is a domain typically found in two-component bacterial sensor and chemotaxis proteins (Schultz *et al.*, 2015) and in eukaryotic histidine kinases (Defosse et al., 2015), and the cyclic nucleotide binding domain found in AAP2. Both, cAMP or cGMP, are key molecules in activating the pathways necessary for egress (Uboldi *et al.*, 2018). The extensive phosphorylation of AAPs could be mediated as well by PKA or PKG to modulate function of the annuli in different stages of the lytic cycle. Furthermore, the Rabaptin-like coiled-coil domain in AAP2 and the GGN-like domain in AAP4 suggest a putative role in vesicular trafficking.

Combining all these insights generates a tentative function of the apical annuli as a (diffusion and/or vesicular trafficking mediated) pore over the IMC during internal budding, and the annuli may serve in addition as a signaling platform during transitions between the intracellular to extracellular state.

## Material and Methods

### Parasites and mammalian cell lines

Transgenic derivatives of the RH strain were maintained in human foreskin fibroblasts (HFF) as previously described (Roos *et al.*, 1994). Parasite transfections and selections were done using 1 μM pyrimethamine, 20 µM chloramphenicol, 20 µM 5’-fluo-2’deoxyuridine (FUDR), or a combination of 25 mg/ml mycophenolic acid and 50 mg/ml xanthine (MPA/X).

### Plasmids and parasite strain generation

*Aap* genes were tagged via endogenous 3’-end replacement. The CAT selection cassette in the tub-YFPYFP(MCS)/sagCAT plasmid (Anderson-White et al., 2011) was replaced with a DHFR/TS selection cassette and the tub-YFP-YFP section was further replaced with a triple-Myc-epitope tag via PmeI and AvrII cloning sites (all oligonucleotide sequences provided in Table S2). Homologous 3’end flanks of a given *aap* gene were cloned via PmeI and AvrII into the generated plasmid. 50 µg of plasmid DNA was linearized with a restriction enzyme digest (Table S3) before transfection in RHΔKu80 parasites.

For endogenous 5’-end tagging of Centrin2 we used a CRISPR-based strategy. Transfection of 40 µg of pU6-Universal plasmid (kindly provided by Sebastian Lourido, Addgene: 52694), carrying a suitable sgRNA (Table S3), resulted in a DNA-double strand break around the ATG of the gene. The break was repaired by co-transfection of 40 µg of a DNA ultramer that had homologous regions at its 5’- and 3’-end, contained a LoxP recombination-site as well as a tandem Myc-epitope tag. Parasites that integrated the ultramer were identified by single cell cloning using limiting dilution and IFA. To generate the cytosolic BioID2-YFP control we cloned the *bioid2* coding sequence (Kim et al., 2016) via BglII and AvrII cloning sites in the tub-YFP-YFP (MCS)/sagCAT plasmid. The tubulin promoter was subsequently replaced with the *morn1* promoter sequence (Gubbels *et al.*, 2006) via a PmeI/BglII restriction sites. For endogenous 5’-end tagging of AAP4 with BioID2 we used a previously reported method that links expression of a selection marker to the integration into a specific gene locus (selection-linked integration (SLI) (Birnbaum et al., 2017)). Hereto we designed a plasmid in which the HXGPRT ORF was followed by the sequence of the 2A-like peptide from *Thosea asigna* virus (T2A; (Szymczak *et al.*, 2004, Straimer *et al.*, 2012)), followed by the coding sequence for the Ty-epitope tag and the *bioid2* coding sequence. All parts were assembled using Gibson assembly strategy (NEB). The established plasmid served as a template for PCR amplification with oligomers that added 35 bp homologous flanks, surrounding the AAP4 start codon, to the resulting PCR product. A DNA-double strand break was made through co-transfection of 40 µg of an AAP4-ATG CRISPR/Cas9 plasmid and parasites were selected using MPA/X for expression of HXGPRT under the endogenous *aap4* promoter.

AAP2-cKD parasites were generated by cloning a homologous 5’-flank of the ORF via BglII and NotI restriction sites into a DHFR-TetO7-sag4-Ty plasmid. This plasmid was linearized with NheI and subsequently transfected in TATiΔKu80 parasites (Sheiner *et al.*, 2011).

AAP4-KO parasites were generated using a CRISPR-based strategy. Two pU6 plasmids (20 µg of DNA for each), carrying sgRNAs directed against the 5’ and 3’ end of the *aap4* ORF, were transfected in RHΔKu80 parasites, together with a DHFR resistance marker cassette that had 35bp homologous overhangs to the 5’- and 3’-UTRs of the gene. For genetic complementation of the AAP4-KO, a plasmid that expresses AAP4-Myc_3_ under its endogenous promoter flanked by homologous regions to the 5’- and 3’-UTR of the *uprt* locus was generated and integrated into the *uprt* locus of AAP4-KO parasites. Pseudodiploid expression of the Ty-BioID2-Centrin2 fusion-gene was generated by cloning the pmorn1-Ty-BioID2 sequence via PmeI and AvrII restriction sites in a previously generated pmorn1-Myc2-Centrin2-DHFR plasmid (Engelberg *et al.*, 2016). To co-localize AAP4 with Ty-tagged Centrin2, a plasmid was used that contains a 2500 bp flank upstream of the Centrin2 ORF, the Ty-tagged Centrin2 ORF flanked by loxP sites and a 2500 bp flank downstream of the Centrin2 stop codon. All parts were cloned into the previously described plasmid for Cre loxP-based recombination (Andenmatten *et al.*, 2012) and 20 μg of plasmid DNA was transfected in AAP4-Myc_3_ expressing parasites.

### BioID sample preparation and mass spectrum analysis

Biotin labeling for Ty-BioID2-Centrin2 and Ty-BioID2-AAP4 cell lines was done in two biological replicates (+biotin condition) and one biological replicate (-biotin condition). Each biological replicate was run as two technical replicates on the mass spectrometer. The cytosolic BioID2-YFP control was done as one technical replicate for the (+) and (-) biotin condition.

Parasites expressing BioID2-fusion proteins were grown overnight ± 150 μM biotin and harvested by mechanical lysis in 1%SDS in resuspension buffer (150 mM NaCl, 50 mM Tris-HCl pH 7.4). For the streptavidin pull-down, 1.5 mg of total protein lysate was cleared by centrifugation and mixed with Streptavidin-agarose beads (Fisher) in 1% SDS in DPBS (Corning). Beads and lysates were incubated overnight at 4°C and proteins bound on beads were washed the next day with 0.1% SDS in DPBS, DPBS and H_2_O. Beads were resuspended in 6 M Urea in DPBS, reduced, alkylated and digested with 2 μg of Trypsin (Promega) overnight at 37°C. Digested peptides were separated from beads by centrifugation and subsequent mass-spectrometric analysis was performed.

LC-MS/MS analysis was performed on an LTQ-Orbitrap Discovery mass spectrometer (ThermoFisher) coupled to an Agilent 1200 series HPLC. Samples were pressure loaded onto a 250 µm fused silica desalting column packed with 4 cm of Aqua C18 reverse phase resin (Phenomenex). Peptides were eluted onto a biphasic column (100 µm fused silica column with a 5 µm tip packed with 10 cm Aqua C18 resin and 4 cm Partisphere strong cation exchange resin (SCX, Whatman)) using a gradient of 5-100% Buffer B in Buffer A (Buffer A: 95% water, 5% acetonitrile, 0.1% formic acid; Buffer B: 20% water, 80% acetonitrile, 0.1% formic acid). Peptides were then eluted from the SCX resin onto the Aqua C18 resin and into the mass spectrometer using four salt steps (95% water, 5% acetonitrile, 0.1% formic acid and 500 mM ammonium acetate) ((Weerapana *et al.*, 2007)). The flow rate through the column was set to ∼0.25 µL/min and the spray voltage was set to 2.75 kV. With dynamic exclusion enabled, one full MS scan (FTMS) (400-1,800 MW) was followed by 7 data dependent scans (ITMS) of the *n*th most abundant ions.

The tandem MS data were searched using the SEQUEST algorithm ((Eng *et al.*, 1994)) using a concatenated target/decoy variant of the *Toxoplasma gondii* GT1 ToxoDB-V29 database. A static modification of +57.02146 on cysteine was specified to account for alkylation by iodoacetamide. SEQUEST output files were filtered using DTASelect 2.0 ((Tabb *et al.*, 2002)). Reported peptides were required to be unique to the assigned protein (minimum of two unique peptides per protein) and discriminant analyses were performed to achieve a peptide false-positive rate below 5%.

### Analysis of mass-spec data by probabilistic calculation of interactions

Spectral counts of unique proteins were used to determine probability of interaction for given bait and preys using SAINTexpress (Teo et al., 2014). SAINTexpress was executed using the –L4 argument, compressing the four largest quantitative control values of a given prey in one virtual control. The resulting SAINTexpress matrix was visualized using the Prohits-viz online suite (Knight *et al.*, 2017) with the following settings: abundance column set to “spectral sum [SpecSum]”, score column set to “average probability [AvgP]”, primary filter = 0.8, secondary filter = 0.6, to generating the dot plot and the prey-prey correlation. The following preys were manually deleted from the analysis using the Zoom function in Prohits-viz: HXGPRT, TGGT1_269600 (annotated as biotin enzyme), TGGT1_289760 (annotated as biotin-synthase). An output of the dot plot analysis with relaxed settings (primary filter: 0.5; secondary filter: 0.3) was used to visualize the interaction network with Cytoscape_v3.6.1 (Koh *et al.*, 2012, Saito *et al.*, 2012, Shannon *et al.*, 2003).

### AAP4 and IMC3 antiserum generation

To generate N-terminal His_6_ tagged fusion protein, 1143 bp from the cDNA of AAP4 (corresponding to amino acids: 121 to 500) were PCR amplified using the primers pAVA0421-AAP4-FR and cloned into the pAVA0421 plasmid (Alexandrov *et al.*, 2004) by Gibson Assembly (NEB). The fusion protein was expressed in BL21 Codon Plus (DE3) RIPL *Escherichia coli* (Agilent) using 0.5 mM IPTG in 2xYT broth 5 hrs at 37°C and purified under native condition over Ni-NTA Agarose (Invitrogen). Polyclonal antiserum was generated by immunization of a guinea pig (Covance, Denver, PA). Antisera were affinity purified as previously described (Gubbels et al., 2006) against recombinant His_6_-AAP4(121-500).

His_6_-TgIMC3(N1-120) was expressed as described before (Anderson-White et al., 2011) and polyclonal serum was generated by immunization of a rabbit (Covance, Denver, PA).

### (Immuno-) fluorescence microscopy

Indirect immunofluorescence assays were performed on intracellular parasites grown overnight in 6-well plate containing coverslips confluent with HFF cells fixed with 100% methanol (unless stated otherwise) using the following primary antisera: mouse α -Ty clone BB2 (1:500); kindly provided by Sebastian Lourido, Whitehead Institute, MAb 9E10 α-cMyc (1:50); Santa Cruz Biotechnology, MAb 9B11 α-cMyc Alexa488 conjugated (1:100); Cell Signaling Technologies, rabbit α-beta tubulin (1:1,000); kindly provided by Naomi Morrissette, University of California, Irvine (Morrissette & Sibley, 2002), rat α-IMC3 (1:2,000 (Anderson-White et al., 2011)), rabbit α-IMC3 (1:2,000), rat α-ISC2 (1:1000); kindly provided by Peter Bradley, University of California, Los Angeles (Chen et al., 2016)) and guinea pig α-AAP4 (1:200). Streptavidin-594 (1:1000); Thermo Fisher, Alexa 488 (A488) or Alexa594 (A594) conjugated goat α-mouse, α-rabbit, α-rat, or α-guinea pig were used as secondary antibodies (1:500); Invitrogen. DNA was stained with 4’,6-diamidino-2-phenylindole (DAPI). A Zeiss Axiovert 200 M wide-field fluorescence microscope was used to collect images, which were deconvolved and adjusted for phase contrast using Volocity software (Improvision/Perkin Elmer). SR-SIM or Zeiss Airyscan imaging was performed on intracellular parasites fixed with 4% PFA in PBS and permeabilized with 0.25% TX-100 or fixed with 100% methanol. Images were acquired with a Zeiss ELYRA S.1 and Airyscan system in the Boston College Imaging Core in consultation with Bret Judson. All images were acquired, analyzed and adjusted using ZEN software and standard settings. Final image analyses were made with FIJI software.

### Western Blot

Parasites for western blots were filtered through 3 μm filters, washed in 1xPBS before lyzed in resuspension buffer (50mM Tris-HCl, pH 7.8, 150 mM NaCl, 1% SDS and 1x protease inhibitor cocktail (Sigma Aldrich)) and incubated at 95°C for 10 min. Parasites at 10 million/lane were loaded and analyzed by SDS-PAGE. Nitrocellulose blots were probed with MAb 9E10 α-cMyc-HRP (1:4000); Santa Cruz Biotechnology, mouse α-Ty clone BB2 (1:5000), guinea pig α-AAP4 (1:1000), mouse α-GFP (1:2000); Roche, or Streptavidin-HRP (1:4000); Thermo Fisher. Secondary antibodies were conjugated to HRP and used in dilutions of 1:10000 (goat α-mouse, Dako) and 1:3000 (goat α-guinea pig, Santa Cruz Biotechnology). Equal loading was detected with MAb 2 28 33 α-beta-tubulin (1:5000); Invitrogen.

### *In vivo* mouse infection studies

Groups of four C57BL/6J mice with a weight between 18-20 g were infected intraperitoneally with 100 or 1,000 tachyzoites of the RHΔKu80 or AAP4-KO and AAP4-KO complemented strains. Following infection, mice were monitored daily for posture, activity level and weight. All animal protocols were approved by the Institutional Animal Care and Use Committee (IACUC) of Boston College.

## Acknowledgements

We thank Bret Judson and the Boston College Imaging Core for infrastructure and support, Emily Stoneburner for technical support, Drs. Bradley, Lourido, Morrissette, and Ward for sharing reagents.

This study was supported by National Science Foundation (NSF) Major Research Instrumentation grant 1626072, NSF Research Experience for Undergraduates (REU) grant 1560200, National Institute of Health grants AI144856, AI110690, AI110638, and AI128136, a Deutsche Forschungsgemeinschaft grant, and an American Heart Association post-doctoral fellowship grant 17POST33670577. The funders had no role in study design, data collection and analysis, decision to publish, or preparation of the manuscript.

