## Supplementary figures and images for "The apical annuli of *Toxoplasma gondii* are composed of coiled-coil and signaling proteins embedded in the IMC sutures"

### Figure S1

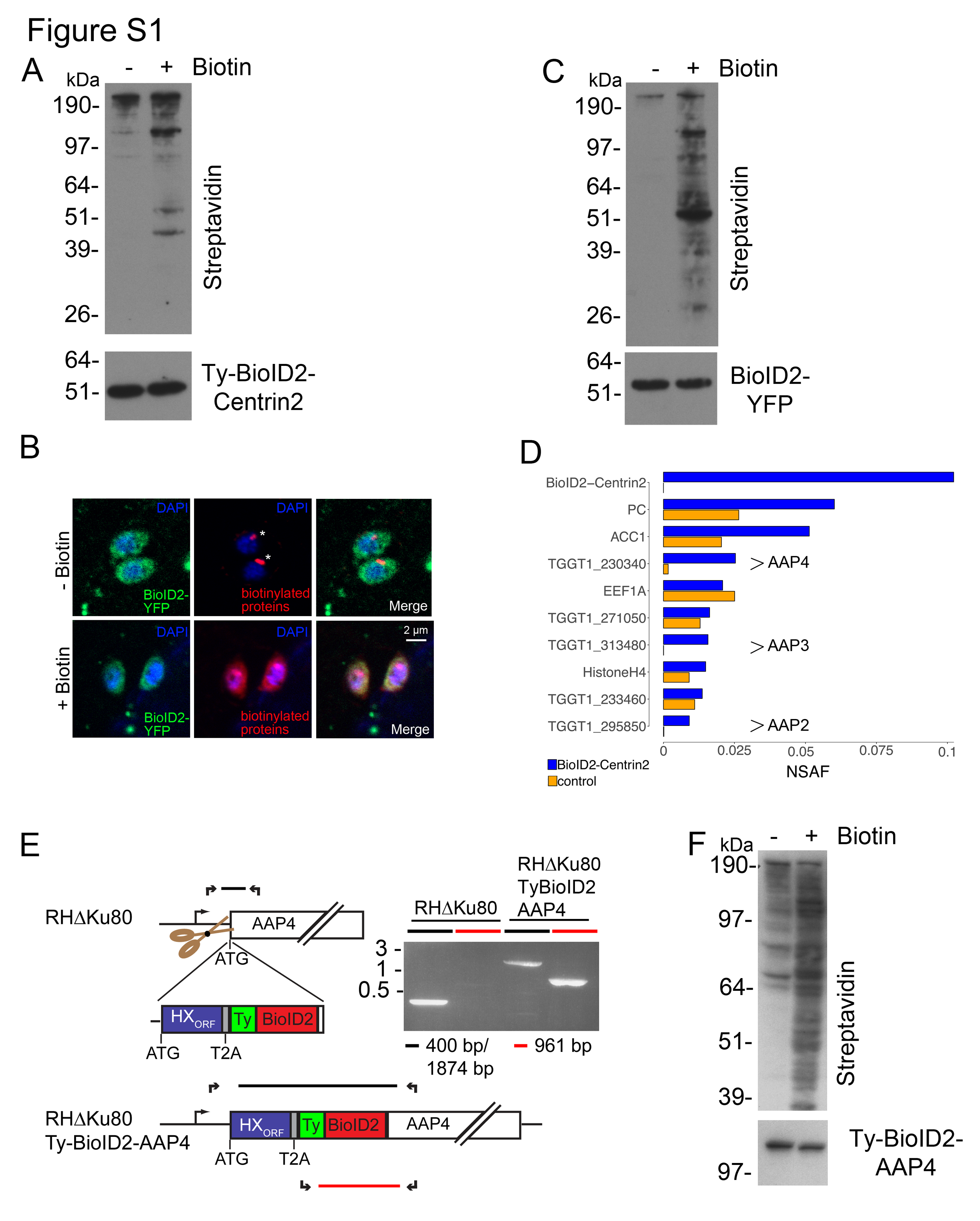

### Figure S2

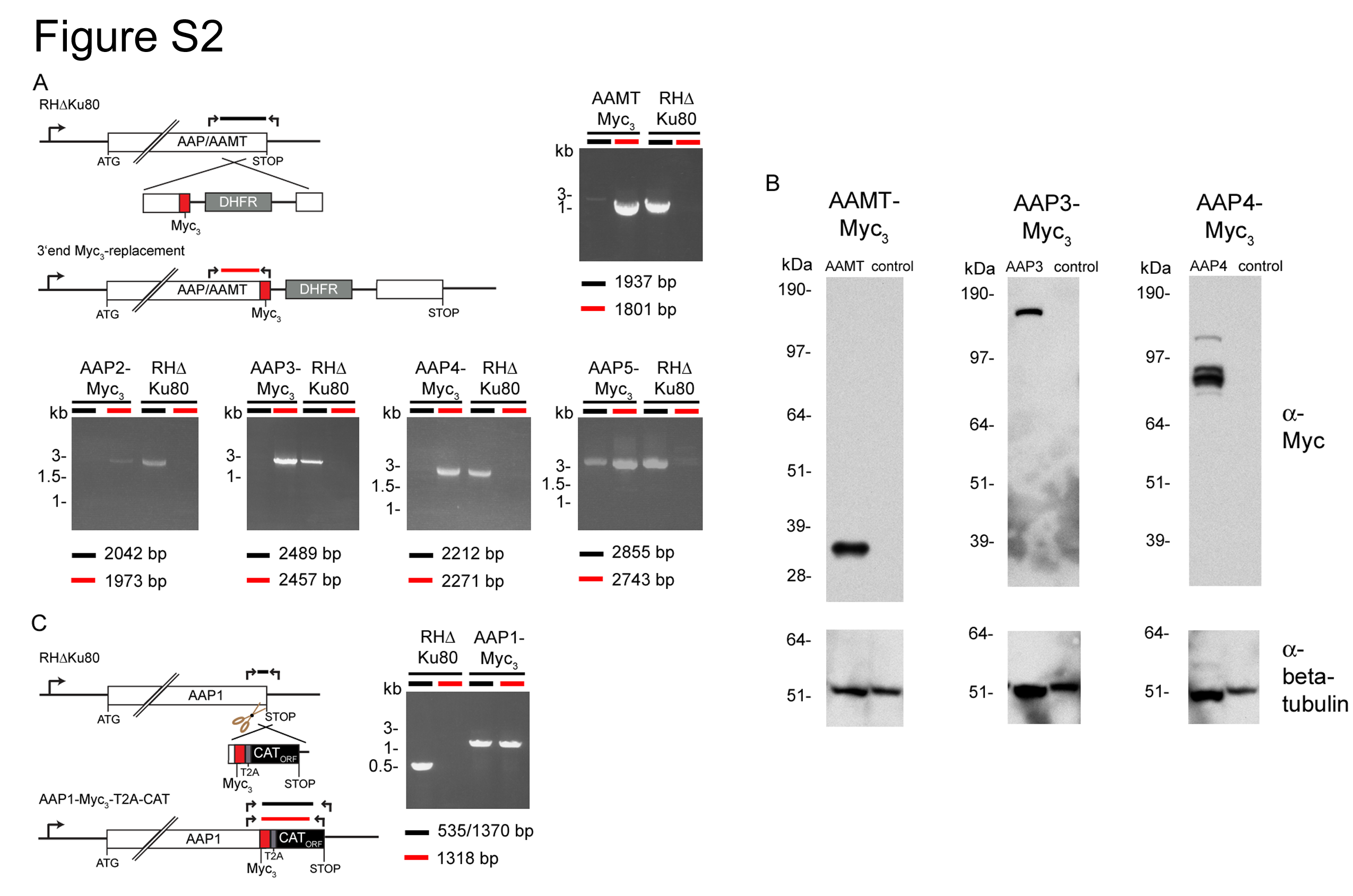

### Figure S3

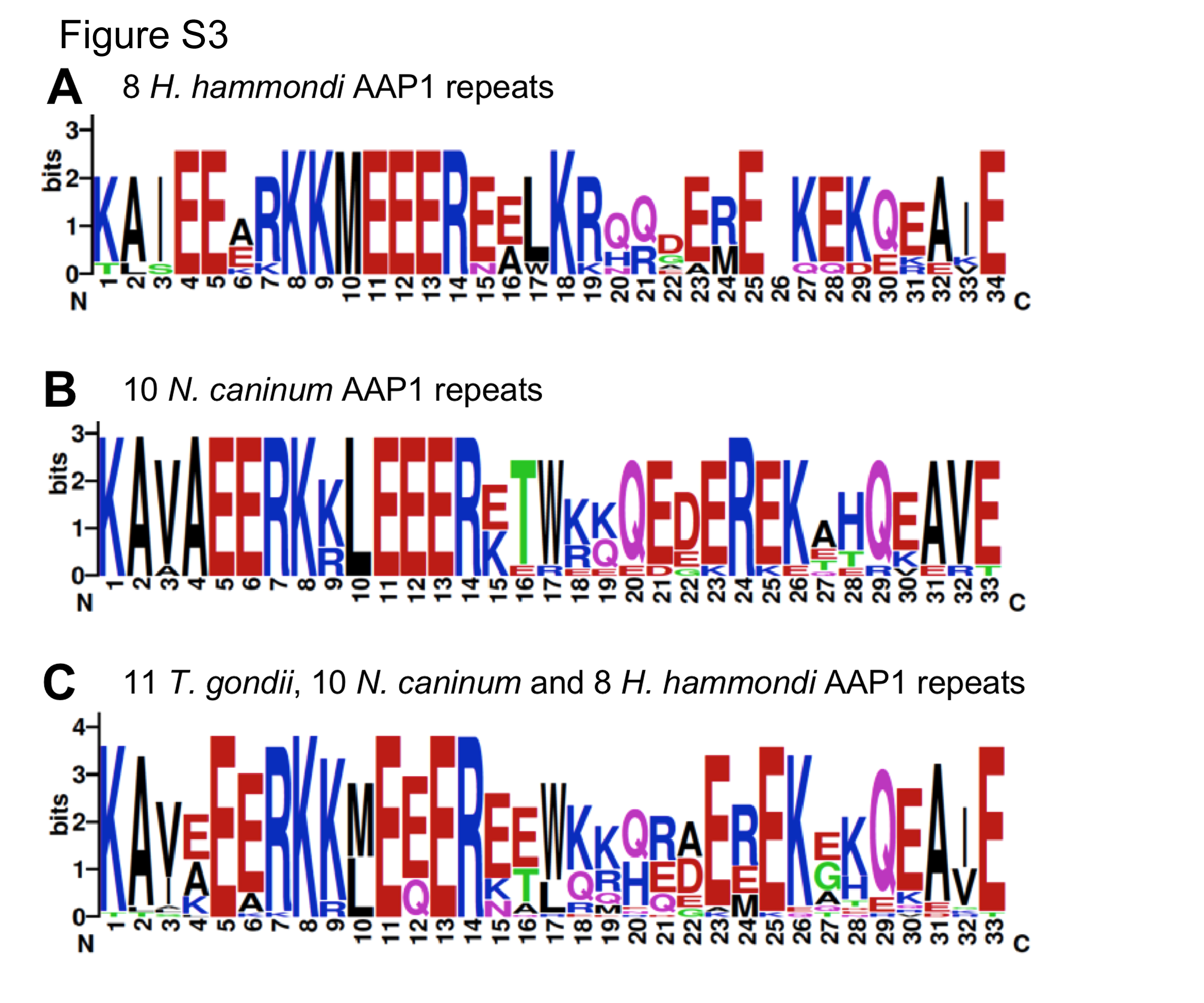

### Figure S4

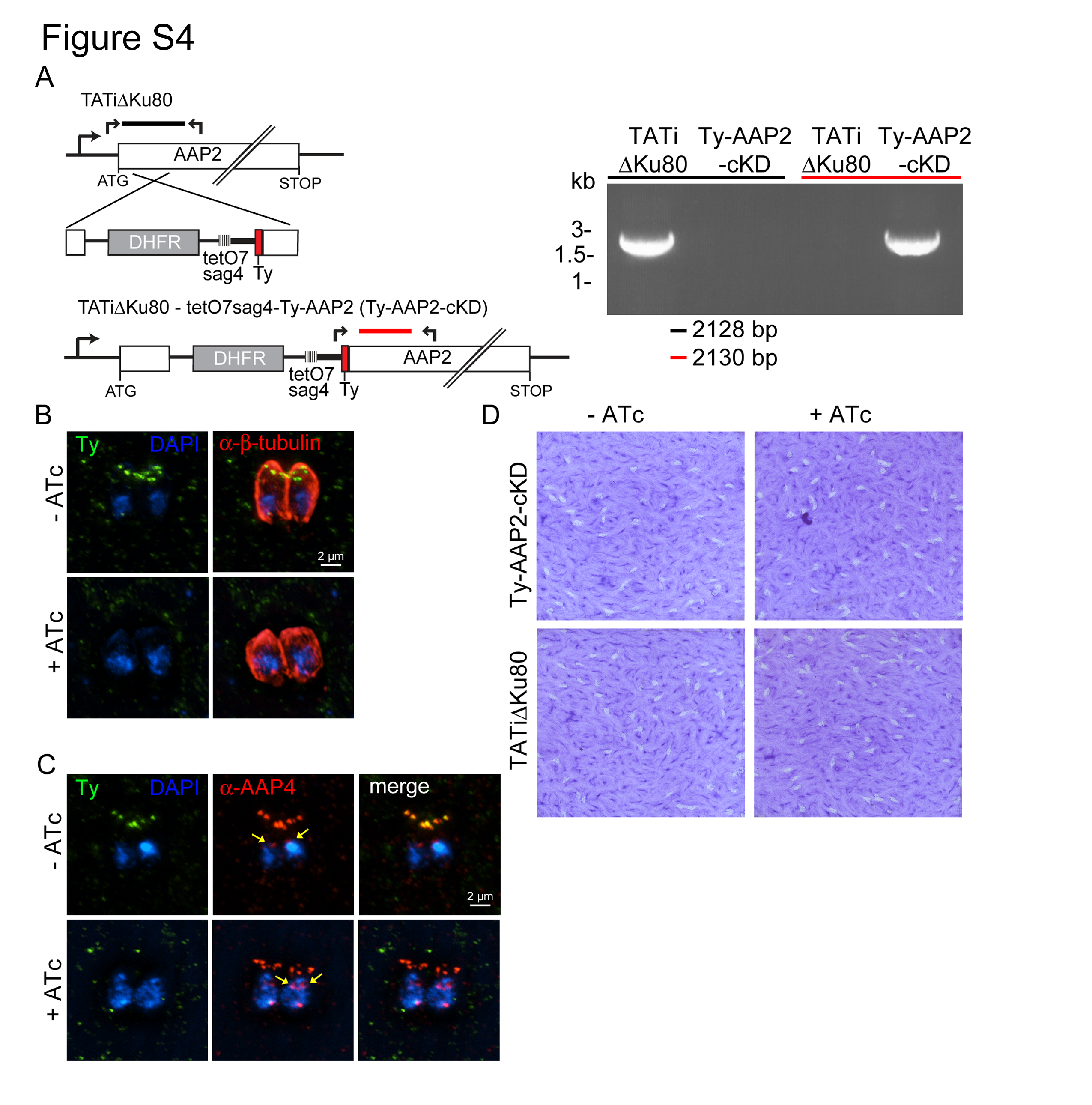

### Figure S5

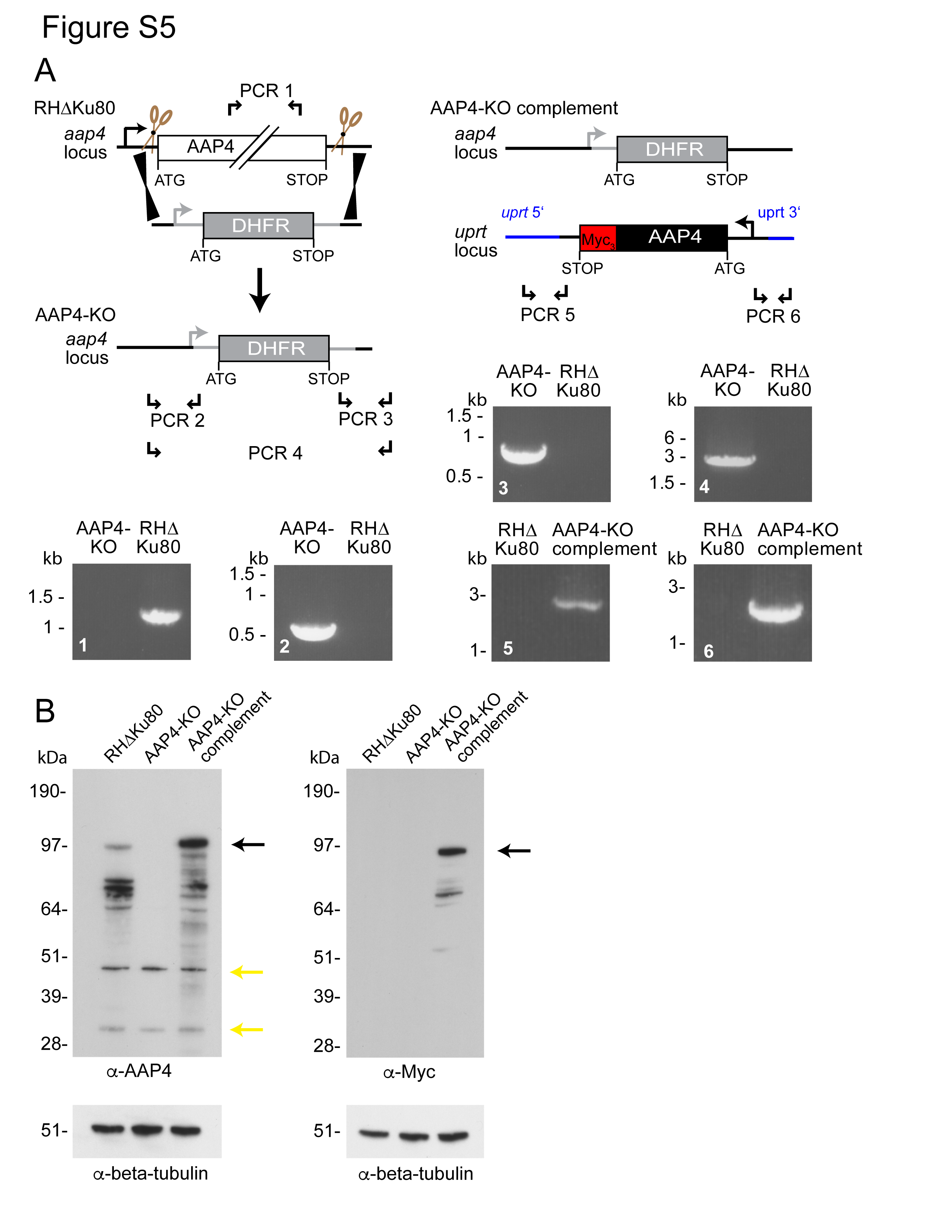

### Figure S6

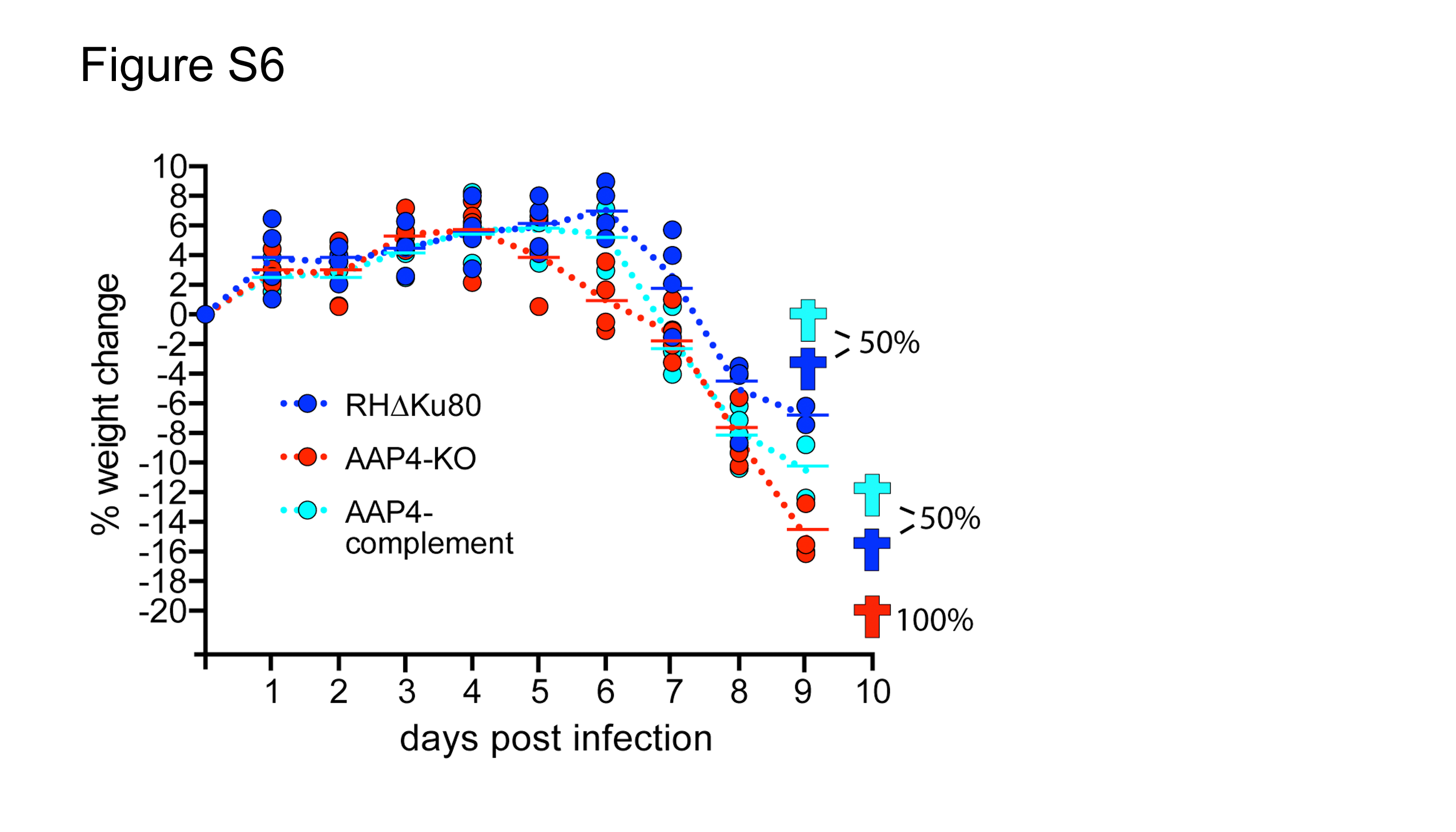
