## Supplementary Table S1 S2 for "The apical annuli of *Toxoplasma gondii* are composed of coiled-coil and signaling proteins embedded in the IMC sutures"

### Supplementary material:

**Table S1. Excel file with SAINTexpress analysis results.**

**Table S2. Oligonucleotides used in this study.** Restriction enzyme sites used for cloning are underlined.

| **AAP-Myc tagging** | **Restriction site** | **Sequence** |
|---|---|---|
| AAP2-Myc_s | PmeI | CAGGTTTAAACCGGCCGACCAGAGACTC |
| AAP2_Myc_as | AvrII | CTGCCTAGGCCGCGGCCCAGCCATC |
| AAP3_Myc_s | PmeI | CAGGTTTAAACAAGCAGGGAAGACCTTTCGC |
| AAP3_Myc_as | AvrII | CAGCCTAGGCGACAAATCAACTTCAACAAGAGC |
| AAP4_Myc_s | PmeI | CAGGTTTAAACAGCTTCTTCTCGAGGAAACC |
| AAP4_Myc_as | AvrII | CAGCCTAGGCTGTTTCTGAAGACTCCGG |
| AAP5_Myc_s | PmeI | GACGTTTAAACGTTTTCTTCCTTTTTGATCGACCG |
| AAP5_Myc_as | AvrII | GTCCCTAGGGCGCCTGACAAATCTGTGGTTT |
| AAMT_s | PmeI | CAGGTTTAAACTTGAGAGATACGAACGTGCAG |
| AAMT_as | AvrII | CTGCCTAGGGTTCGTCGGGGCACCTTTG |
| **AAP1-3xMyc-T2A-CAT** | | |
| pU6-AAP13end_s | | AAGTTGGAGCGAAGAAGGACTACAG G |
| pU6-AAP1_3end_as | | AAAAC CTGTAGTCCTTCTTCGCTCC A |
| AAP1-Myc-s | | CCTCGACGCAGACTCGTCGCCTGCGCCTCTGCCTAGGGAACAAAAGTTGATTTCTGAAG |
| AAP1-Myc-as | | GATGTCCGCGCCGCGAACGACCGCGGAGCGAAGAA TTAATTAAGCCCCGCCCTGCCACTC |
| Myc-s | | GGGAACAAAAGCTGGGTACC GAACAAAAGTTGATTTCTGAAGA |
| Myc-T2A-as | | CGTCACCGCATGTTAGCAGACTTCCTCTGCCCTCTCCCAGGTCCTCCTCGGAGATCAGC |
| CAT-T2A_s | | CTAACATGCGGTGACGTCGAGGAGAATCCTGGACCT CATGAGAAAAAAATCACTGGATATACC |
| CAT_as | | TGGAGCTCCACCGCGGTGGCTTAATTAAGCCCCGCCCTGCCACTC |

| | | |
|---|---|---|
| **pmorn1-TyBioID2-Centrin2-DHFR** | | |
| TyBioid2_s | BglII | GACAGATCTAAAATGGAGGTCCATACTAACCAAGATCCACTTGACTTCAAGAACCTGATCTGGCTG |
| Bioid2_as | AvrII | GACCCTAGGGCTTCTTCTCAGGCTGAACTC |
| **AAP4_BioID2** | | |
| AAP4_BioID2_s (binds in HXGPRT) | | ACTCCCTCCCCTCGCGTTCTTCTCCGTTTAAAATGGCGTCCAAACCCATTG |
| AAP4_BioiD2_as (binds at the end of BioID2) | | GGAATCGGAGGAGGCTGGTGGCGAACGGCTTCTTCTCAGGCTGAACTCGCC |
| **HX-T2A-Ty-BioID2** | | |
| HX_s | | GGGAACAAAAGCTGGGTACCAAAATGGCGTCCAAACCCATTGAA |
| HX_T2A_as | | CGTCACCGCATGTTAGCAGACTTCCTCTGCCCTCTCCCTTCTCGAACTTTTTGCGAGC |
| T2A-Ty-BioID2_s | | CTAACATGCGGTGACGTCGAGGAGAATCCTGGACCTGAGGTCCATACTAACCAAGATCCAC |
| BioID2_as | | TGGAGCTCCACCGCGGTGGCGCTTCTTCTCAGGCTGAACTCGCC |
| AAP4_inte_s | | TCAAGA ACTCTCCAACGCG |
| AAP4_inte_as | | AGGAAGCCTGAAGCGACC |
| **BioID2-YFP** | | |
| BioID2_s | BglII | CAGAGATCTAAAATGTTCAAGAACCTGATCTGGC |
| BioID2_as | AvrII | CTGCCTAGGGCTTCTTCTCAGGCTGAACTC |
| **Myc2Centrin2** | | |
| pU6-Cen2_ATG_s | | AAGTT GCGTGTTCCAACATGCAGC G |
| pU6-Cen2_ATG_as | | AAAAC CGCTGCATGTTGGAACACGC A |
| loxP-Myc-Cen2_s | | CCTTCTACATTTGGCCTTTTTTTGTCGAGCGTGTTATAACTTCGTATAGCATACATTA TACGAAGTTATAAAATGGAACAAAAGCTAATCTCCGAGGAAGACTTGAACGGTGCTA GGGCCGAG |

| | | |
|---|---|---|
| loxP-Myc-Cen2_as | | GCTGTCGGGCTCGCCCCTCGCAGTGCTCCTCGCTGCAGGTCCTCCTCGGAGATCAG CTTCTGCTCCTCGGCCCTAGCACCGTTCAAGTCTTCCTCGGAGATTAGCTTTTGTTCCA TTTTATAACTTCGTATAATGTATGCTATACGAAGTTATAACACGCTCGACAAAAAAAGGCC AAATGTAGAAGG |
| **AAP-integration PCR** | | |
| AAP2_inte_s | | TTTGGCAGGAAAGGCCCGC |
| AAP2_inte_as | | ACCGAAAGATGCGTCGTCGC |
| AAP3_inte_s | | ATTCTTGGCGCTTGCATGTGG |
| AAP3_inte_as | | ACGAGATGTGTTCCTCGGAC |
| AAP4_inte_s | | AAAGACAAA CAGCTAGCAGCC |
| AAP4_inte_as | | ATGCTTATGCATGCCTGGTGG |
| AAP5_inte_s | | TTCAACCTAGGGTCGTCT GC |
| AAP5_inte_as | | TCGGATTATCTAAAGTACGGGC |
| AAMT_inte_s | | TTGCGTTTTTGTATGTGGAGGTG |
| AAMT_inte_as | | AGAGCAATCTGTGACTACGGC |
| Myc_as | | TCAGCTCTCTCCTTATTACAGG |
| **AAP4-KO** | | |
| pU6-AAP4_5'end_s | | AAGTT GGCTGGTGGCGAACGCATAG G |
| pU6-AAP4_5'end_as | | AAAAC CTATGCGTTCGCCACCAGCC A |
| pU6-AAP4_3'end_s | | AAGTT G TTCACCTCTGGTGTATCCCT G |
| pU6-AAP4_3'end_as | | AAAAC AGGGATACACCAGAGGTGAAC A |
| AAP4_5end-DHFR_s | | TTCTCCACTCCCTCCCCTCGCGTTCTTCTCCGTTT CACGAAACCTTGCATTCAAACC |
| AAP4_3end_DHFR | | CGCCTGCGCGGGGGCGAAGGTCTGCGGCGAGACTC ATCCTGCAAGTGCATAGAAGG |
| AAP4_ORF_inte_check_s | | TCTCGTCTCTGTCTCAAGGG |
| AAP4_ORF_inte_check_as | | ATGTCTCGTCGCTCTTCTCG |

| | | |
|---|---|---|
| AAP4_complementation_promoter_s | | CAGCCTCACGTTAACGCGGCCGCCGTTTAAACGCACTTCTTGGCGCAGATCCG |
| AAP4_complementation_promoter_as | | TGGTGGCGAACGCATTTTTTAATTAAAGAGGAAACGGAGAAGAACGCG |
| AAP4_ORF_s | | TTCTTCTCCGTTTCCTCTTTAATTAAAAAATGCGTTCGCCACCAGCCTC |
| AAP4_ORF_as | | TTCTTCAGAAATCAACTTTTGTTCGCTAGCCTGTTTCTGAAGACTCCGGAG |
| UPRT_s (verifies KO of UPRT) | | CGTTTCTTTACTGGCATCGAATG |
| UPRT_as (verifies KO of UPRT) | | GTTGTTTCGTCTCTCTGGATG |
| UPRT_5end_s | | CGGTGTGGTTCCTGTTGACTTAG |
| UPRT_3end_as | | GTGCAGGGAGGTTTGTTATCTTG |
| AAP4_promoter_as (to verify integration in UPRT locus | | AACAGGTACGCAAGACTGTCG |
| **AAP4- His-expression** | | |
| pAVA0421-AAP4-F | | GAAGCTCAGACCCAGGGTCCTGGTTCGATGCACAGCGACTTCGTTTCGACG |
| pAVA0421-AAP4-R | | TGCAGAACTTGTTCGTGCTGTTTTATGCTAGCTGTTTGTCTTTCTGCAG |
| **TetO7-sag4-Ty-AAP2** | | |
| AAP2_s | BglII | CAGAGATCTACCTCCCCCTCGTCTCCGC |
| AAP2_as | NotI | CAGGCGGCCGCAAGCCGCGAAGGTTGGAAGG |
| AAP2_integration_check5'_s | | CTTCCGTAGTTTCTTCTGTTCC |
| AAP2_integration_check3'_as | | CTTCCTGGAAAAAGATGTGGC |
| Sag4_s | | AGCATACTGCAACTGCTTTCG |
| **cen2_5'-loxP-Ty-Centrin2-loxP-YFP-cen2_3'** | | |
| Cen2 5'flank_s | KpnI | GACGGTACCCGTAAGCTCATGCATCGC |
| Cen2_5'flank_loxP_as | EcoRI | GACGAATTCATAACTTCGTATAATGTATGCTATACGAAGTTATGGAACACGCTCGACAAAAAAAGGCC |
| Cen2_ORF_s | EcoRI | GACGAATTCAAAATGCAGCGAGGAGC |
| Cen2_ORF_as | PacI | GACTTAATTAACTACGGGAAAGTCTTCTTGG |

| | | |
|---|---|---|
| Cen2_3’flank_s | SacI | GACGAGCTCCAAGGCTGTCGATTCAACAGAGAG |
| Cen2_3’flank_as | SacI | GACGAGCTCAAACAGCGATTCTCAGAGACGC |

### Table S3. Plasmids generated for this study

| Name | Reference | Restistance marker | Unique restriction site | sgRNA sequence | Description |
|---|---|---|---|---|---|
| AAP2-Myc$_3$ | this paper | DHFR | NsiI | – | For 3'end tagging of AAP2 |
| AAP3-Myc$_3$ | this paper | DHFR | SfiI | – | For 3'end tagging of AAP3 |
| AAP4-Myc$_3$ | this paper | DHFR | NsiI | – | For 3'end tagging of AAP4 |
| AAP5-Myc$_3$ | this paper | DHFR | XmaI | – | For 3'end tagging of AAP5 |
| AAMT-Myc$_3$ | this paper | DHFR | MfeI | – | For 3'end tagging of AAMT |
| pmorn1-BioID2-YFP | this paper | CAT | – | – | For control BioID2 experiments |
| pmorn1-TyBioID2-Centrin2 | this paper | DHFR | – | – | For Centrin2 biotinylation experiments |
| pU6-Cas9-Universal | Addgene:52694 | – | – | – | Backbone for pU6-Cas9 constructs |
| pU6-Cas9-AAP4-ATG | this paper | – | – | GGCTGGTGGCGAACGCATAG | For 5' integration in AAP4 locus |
| pU6-Cas9-AAP4-STOP | this paper | – | – | GTTCACCTCTGGTGTATCCCT | For 3' integration in AAP4 locus |
| pU6-Cas9-AAP1-STOP | this paper | – | – | GGAGCGAAGAAGGACTACA | For 3' integration in the AAP1 locus |
| pU6-Cas9-Centrin2-ATG | this paper | – | – | TGCGTGTTCCAACATGCAGC | For integration in the 5' Centrin2 locus |
| tetO7-sag4-Ty-AAP2 | this paper | DHFR | NheI | – | For promoter replacement of AAP2 |
| Myc$_3$-T2A-CAT$_{ORF}$ | this paper | CAT | – | – | As template for AAP1 3'integration |
| HXGPRT$_{ORF}$-T2A-Ty-BioID2 | this paper | HXGPRT | – | – | As template for 5' BioID2 integration in the AAP4 locus |
| cen2_5’-loxP-Ty-Centrin2-loxP-YFP-cen2_3’ | this paper | HXGPRT | KpnI | – | To co-localize AAP4-Myc$_3$ with TyCentrin2 |
