## Supplementary material for "The apical annuli of *Toxoplasma gondii* are composed of coiled-coil and signaling proteins embedded in the IMC sutures": Suplementary Figure and Video Legends

**Supplementary Figure and Video legends**

**Figure S1. Validation of biotinylation capacities of utilized BioID2 fusion proteins, identification of AAP4 and construction of Ty-BioID2-AAP4.**

**A. C. F.** Western blots of total parasite lysates using streptavidin-HRP displaying increased protein biotinylation after BioID activation. Parasites were grown overnight in presence of ±150 μM biotin. Fusion proteins were detected with α-Ty (**A:** Ty-BioID2-Centrin2 and **F:** Ty-BioID2-AAP4) or α-GFP antisera (**C:** BioID2-YFP), which were also used as loading controls. **B.** IFAs of the cytosolic BioID2 control. BioID-YFP fusion protein is indicated in green, co-stained with streptavidin-A594 (red) and DAPI (blue). Parasites were incubated ±150 μM biotin overnight before IFA. Enriched biotinylation is detected in presence of biotin in the cytosol of the parasite. Note that streptavidin-A594 at all times highlights to the apicoplast (asterisks), which naturally harbors biotinylated proteins. **D.** Top 10 identified proteins ordered (top = highest) according their NSAF in the Centrin2 data set. Comparison of the calculated NSAF of the BioID2-Centrin2 (blue) and the cytosolic control (orange) data set is shown. TGGT1_230340: AAP4; TGGT1_313480: AAP3 and TGGT1_295850: AAP2 show specific enrichment in the BioID2-Centrin2 data set. PC: pyruvate carboxylase, TGGT1_284190; ACC1: acetyl-CoA carboxylase, TGGT1_221320; EEFA1: elongation factor 1 alpha, TGGT1_286420A; Histone H4: TGGT1_239260. **E.** Generation of endogenously N-terminally Ty-BioID2-tagged AAP4. The endogenous AAP4 promoter drives the HXGPRT selectable marker that is fused through a T2A skip peptide to the Ty-BioD2-AAP4 open reading frame (Birnbaum et al., 2017). Integration was facilitated by a co-transfected CRISPR/Cas9 plasmid that targets the 5’end of the gene (scissors). Right panel shows PCR validation of the genotype.

**Figure S2. The AAP1 repeats in *Hammondia* *hammondi* and *Neospora caninum.* A.** Logo plot of the eight HhAAP1 repeats. The empty spot at position 26 derives from a K residue insertion only present in repeat number 5. **B.** Logo plot of the ten NcAAP1 repeats. **C.** Logo plot combining all AAP1 repeats present in *Toxoplasma*, *Hammondia* and *Neospora*. For this plot, the K residue was manually removed from repeat 5 in HhAAP1.

**Figure S3. Verification of generated AAP parasite cell lines. A.** Endogenously tagged AAP2-AAP5, and AAMT lines were generated by transfection of an integration plasmid, which harbors a crossover flank to target the 3’-end of the respective gene. Oligonucleotides that either bind in the endogenous 3’-UTR (black bars) of the *aap* genes or in the Myc-tag sequence (red bars), were used to verify integration events by PCR. B. Western blot analysis of endogenously tagged AAP and AAMT proteins. We detected the predicted sizes for the AAP proteins 3 and 4 (152 and 102kDa, respectively) and for the methyltransferase AAMT (24 kDa) with a specific anti-Myc antibody. We were unable to detect protein bands for the AAP proteins 2 and 5, presumably due to their predicted high molecular weight (AAP2: 331 kDa and AAP5: 265 kDa). Equal loading was confirmed with anti-beta-tubulin (clone 2 28 33) antibody. **C.** To target the endogenous 3’end of *aap1* with a triple Myc tag, we used the previously described SLI approach. Hereto, we PCR amplified a triple Myc-tag fused via a T2A skip peptide to the Chloramphenicole acetyltransferase (CAT) ORF and transfected it together with a CRISPR/Cas9 plasmids that targets the 3’-end of the gene (scissors). PCR verified correct integration in the established AAP1-Myc_3_ line.

**Figure S4. Generation and characterization of AAP2 conditional knockdown (cKD) parasites. A.** To generate AAP2cKD parasites, we replaced the endogenous 5’-end of the genes with a version that is driven by the regulatable TetO7sag4 promoter. Integration was achieved through single crossover in the 5’-end of the *aap2* ORF. PCR verified the correct integration. **B.** This regulatable line exhibits AAP2 signal at the apical annuli, as observed with Ty-antiserum (green). Upon gene knockdown (+ATc) the AAP2 signal is no longer observed in the parasite. Periphery of the parasite is visualized with Tg-β-tubulin antiserum staining (red). **C.** Ty-tagged AAP2 co-localizes with AAP4 (red) in the parasite, but AAP4 abundance or localization is not affected upon AAP2 knockdown. Yellow arrows indicate cross-reactive signal seen with the AAP4 antiserum close to the nucleus. Blue: DAPI. **D.** AAP2cKD parasites were grown for 7 days to observe phenotypic consequence of AAP2 knockdown. Plaque number and size did not show significant differences between AAP2cKD parasites that were grown ± ATc or control (TATi∆Ku80) parasites.

**Figure S4. Generation and characterization of AAP2 conditional knockdown (cKD) parasites. A.** To generate AAP2cKD parasites, we replaced the endogenous 5’-end of the genes with a version that is driven by the regulatable TetO7sag4 promoter. Integration was achieved through single crossover in the 5’-end of the *aap2* ORF. PCR verified the correct integration. **B.** This regulatable line exhibits AAP2 signal at the apical annuli, as observed with Ty-antiserum (green). Upon gene knockdown (+ATc) the AAP2 signal is no longer observed in the parasite. Periphery of the parasite is visualized with Tg-β-tubulin antiserum staining (red). **C.** Ty-tagged AAP2 co-localizes with AAP4 (red) in the parasite, but AAP4 abundance or localization is not affected upon AAP2 knockdown. Yellow arrows indicate cross-reactive signal seen with the AAP4 antiserum close to the nucleus. Blue: DAPI. **D.** AAP2cKD parasites were grown for 7 days to observe phenotypic consequence of AAP2 knockdown. Plaque number and size did not show significant differences between AAP2cKD parasites that were grown ± ATc or control (TATi∆Ku80) parasites.

**Figure S6.** **Mouse infection experiment with 100 parasites as inoculum.** Four C57BL/6J mice per group were infected with 100 control, AAP4-KO or AAP4-KO complement parasites on day 0. Weight changes were monitored and plotted in relation to the starting day. While 50% of the mice in the control and AAP4-KO complement group died at day 9, all mice in the AAP4-KO group survived until day 10. Weight change patterns did not show significant differences between the groups, as tested by one-way ANOVA test. Horizontal bars represent the group average; round symbols represent the weight changes of an individual mouse.

**Movie S1 and S2.** 3D-reconstructed SR-SIM image of parasites expressing endogenously tagged AAP4-Myc_3_ co-stained with β-tubulin serum rotated around the y-axis (movie S1) and x-axis (movie S2) corresponding to Figure 5A.

**Movie S3.** 3D-reconstructed SR-SIM image of parasites expressing endogenously tagged AAP4-Myc_3_ co-stained with ISC2 serum corresponding to Figure 5D. Note that ISC2 resides at the apical end of the longitudinal sutures where they meet the apical cap boundary.
